# Emergence of mutants in bacterial populations: a simulation-based approach for parameter inference in extended Luria & Delbrück scenarios

**DOI:** 10.64898/2025.12.19.695396

**Authors:** Aurélien Tauzin, Antoine Frenoy

## Abstract

Bacterial mutation rates are traditionally inferred from phenotypic data using fluctuation assays. Mutation rate computation from these assays relies on a mathematical model describing the emergence of mutants during population growth. However, this standard model relies on restrictive assumptions that are often transgressed in real biological setups. Key assumptions include equal growth rate of the wild-type and the mutant population, no cell death during the growth phase, and no post-growth sampling. Although several refined mathematical models have been proposed over the last decades to circumvent some of these assumptions, none can fully account for the complex ecological scenarios frequently observed in the lab or in the wild, such as the combination of cell death and an unknown fitness effect of the mutation, or non-constant death rates. In this work, we propose to replace these standard mathematical models by a stochastic simulation framework, which can circumvent all these assumptions. We provide a fast and accurate implementation of this stochastic model, and combine it with an Approximate Bayesian Computation scheme to perform simulation-based, likelihood-free parameter inference. Our approach can simultaneously infer mutation rate and fitness effect of the mutation while accounting for death and partial plating, and could be extended to arbitrary complex conditions that can be efficiently simulated.

**Teaser text:** We tackle the challenge of estimating bacterial mutation rates in arbitrarily complex conditions where 3 existing methods do not work (complex demographics, fitness effect of the mutation, etc). We propose a 4 simulation-based method, emphasizing the power of such tools to improve our understanding of mecha5 nisms at the heart of evolutionary processes.

## 1 Introduction

Mutations are the basis of the adaptation and evolution for every living organism. These errors happening during DNA replication or repair, causing a modification of the genetic information, are mostly neutral or deleterious, leading to a dysfunction of a stable biological system (Kimura, 1985). However, mutations are essential for the evolutionary process as they can trigger the emergence of new capacities increasing reproduction and survival. Therefore, the mutation rate is a central parameter to understand evolutionary dynamics. It is a complex trait, reported to be influenced by several genetic and environmental factors. These include DNA repair and more generally DNA replication fidelity systems (Fijalkowska et al., 2012), nutritional stress (Maharjan & Ferenci, 2017, 2018), population density (Krašsovec et al., 2018), and temperature (Van Eldijk et al., 2024). Additionally, these genetic and environmental factors can act jointly on the mutation rate, as it is the case with error-prone repair mechanisms (Goodman, 2002; MacLean et al., 2013).

In 1943, Luria and Delbrü ck published their seminal study (Luria & Delbrück, 1943) establishing the spontaneous nature of mutations, which additionally provided a simple experimental method to estimate the mutation rate towards a specific selectable phenotype (T1 phage resistance, in their case). This experiment, named the Luria-Delbrü ck fluctuation test, fueled an interest in evolutionary microbiology, leading to studies with broad scientific questions regarding micro-organisms demography and adaptation, extended to different biological models (Demerec, 1945, 1948; Demerec & Fano, 1945; Oakberg & Luria, 1947; Stocker, 1949). During the following decades, a number of improved methods emerged to evaluate the mutation rate (*µ*) of a bacterial strain using fluctuation tests, with the most widely used in the literature being the Ma Sandri Sarkar maximum likelihood estimator (Ma et al. (1992), further referred to as MSS-MLE), based on the work of Lea and Coulson (1949) who computed the expected distribution of the number of mutants in the fluctuation assay.

However, because they inherently rely on an analytical model of bacterial growth with mutations, these methods come with a list of assumptions regarding the experimental conditions, the cells demography, and the mutagenesis process. These assumptions are presented in Box 1, directly adapted from Foster (2006). This list is non-exhaustive.

**List of assumptions for mutation rate estimation under analytical models**

**Assumptions that we do not intend to contest or cannot circumvent**

1. The initial number of cells is negligible compared to the final number of cells;
2. The proportion of mutants is always small;
3. Reverse mutations are negligible;
4. The probability of mutation is constant per cell-lifetime;
5. The probability of mutation is not influenced by previous mutational events (see Vázquez-Mendoza et al. (2024) for a mathematical treatment of this problem); **Assumptions which can be circumvented with our method (and others)**
6. The growth rates of mutants and non-mutants are the same;
7. Cell death is negligible
8. All mutants are detected; **Assumptions which could be circumvented with adaptations of our method**
9. The cells are growing exponentially;
10. The probability of mutation per cell-lifetime does not vary during the growth of the culture (see Green et al. (2024) and Krašovec et al. (2018) for biological arguments against this assumption);
11. No mutants arise after selection is imposed.

Box 1: **List of classical biological assumptions** made to analyze fluctuation tests data, adapted from Foster (2006). We reorganized these assumptions in three categories : (1) assumptions that we do not challenge or can not circumvent, (2) assumptions that can be circumvented with the simulation-based method presented in this work, as well as some of the other tools discussed, (3) assumptions which could be circumvented with adaptation of our simulation-based method that are not yet implemented.

These simplifications limit the range of biological scenarios to which these methods can be applied. Complex growth conditions frequently encountered in the lab or in nature are sometimes not compatible with these assumptions. Determining the effect of specific environmental conditions on bacterial mutation rates has, for example, triggered important interest, as it impacts the evolutionary response to treatments or environmental changes, but these growth conditions are incompatible with some of the hypotheses listed in Box 1. These complex conditions include (a) Growth in stressed conditions, including nutrient limitation, drug treatments, physical (UV) or chemical (H2O2) stresses (Carvajal-Garcia et al., 2023; Ferenci, 2019; Kohanski et al., 2010; Maharjan & Ferenci, 2017; Shibai et al., 2017; Torres-Barceló et al., 2013), (b) Experimental conditions where resistance can be conferred by pre-exposure and post-exposure mutations (Cirz et al., 2005; Ford et al., 2013), (c) Mutations which impact fitness (for example mutations conferring rifampicin resistance in the frequently studied *rpoB* gene which are not always neutrals, Reynolds (2000) and Vogwill et al. (2016)), (d) Phenotypic or genotypic heterogeneity in the population beyond the mutation of interest (as it is the case with persistence, Gefen and Balaban (2009), and heterogeneously expressed stress responses, Lansch-Justen et al. (2024)).

Analyzing the outcome of fluctuation tests in these non-standard conditions using tools based on the aforementioned assumptions (Box 1) could lead to erroneous results. This problem led to the development of tools relying on more advanced mathematical methods to compute the mutation rate in some of these complex conditions. For example Flan (Mazoyer et al., 2017; Ycart, 2014), rSalvador (Zheng, 2017) and mlemur (Łazowski, 2023) incorporate the effect of different growth rate (fitness) between wild type and mutant cells, as well as partial plating (only a portion of the cultures plated on selective medium).

Additionally, mlemur can take into account constant cell mortality, and Flan can take into account constant mortality of the mutant (but not of the wild-type). Mlemur can take into account phenotypic lag, as proposed by Sun et al. (2018) and Carballo-Pacheco et al. (2020). Flan and mlemur can take into account variation of the final number of cells in parallel cultures, as proposed by Ycart and Veziris (2014).

While these tools circumvent some of the hypotheses mentioned in Box 1, none of the existing tools can work in more complex cases such as situations where (i) cell mortality has to be taken into account alongside an unknown fitness effect of the mutation, or (ii) the growth period displays several phases with different death rates (as observed for growth under stress).

No analytical formula for the expected distribution of the number of mutants exist for these complex conditions, but simulation-based inference methods could be a way to overcome this limitation: effective Approximate Bayesian Computation methods permit likelihood-free parameter inference based on a simulator of the process of interest (Beaumont, 2010, 2019; Csilléry et al., 2010). Following this idea, Lu et al. (2023) for example proposed simulation based-methods to estimate the mutation rate in the specific case of time-varying (piece-wise constant) mutation rate (but only considered a simple scenario in which the simulator can effectively be replaced by a surrogate model). Beyond parameter estimation, other authors also used simulations of the fluctuation assay to understand the effect of alternative scenarios on the results of a fluctuation test (Carballo-Pacheco et al., 2020; Sun et al., 2018). Generally speaking, simulated models can extend the range of biological scenarios which can be considered beyond what mathematical models permit.

In the present work, we (i) propose a novel simulation-based approach, involving an ABC-MCMC algorithm to infer the mutation rate of bacteria growing in arbitrary complex conditions, taking into account death rate and relative fitness, as well as any other factor that can be incorporated in the simulations (eg. plating efficiency), (ii) test it in a broad range of biological scenarios, and compare it with the three frequently used modern methods (rSalvador, Flan, mlemur) to better understand strengths and weaknesses of each tool. As for now, our inference method includes mutation rate estimation while taking into account the fitness effect of the mutation, cell death, and partial plating; as well as multi-phase growth dynamics (piecewise constant cell death). Moreover, the fitness effect of the mutation can also be jointly estimated alongside the mutation rate in all these conditions.

## 2 Development of a fast and accurate forward simulator for the extended fluctuation test

The scenario we consider in the present work, termed Birth-Death-Mutation-Selection-Sampling, is an extension of the classical fluctuation test model, able to consider several additional biological phenomena: fitness effect of the mutation (parameter *f* describing the fitness of the mutant relative to the wild-type), death (of the wild-type and of the mutant cells, parameter *d*, describing the death rate relative to the birth rate), and sampling (partial or imperfect plating, parameter *p* describing the fraction of the mutant population which is detected in the selective assay). This forward model is depicted in Figure 1 and further described in Appendix (section 5.1).

**Figure 1:**
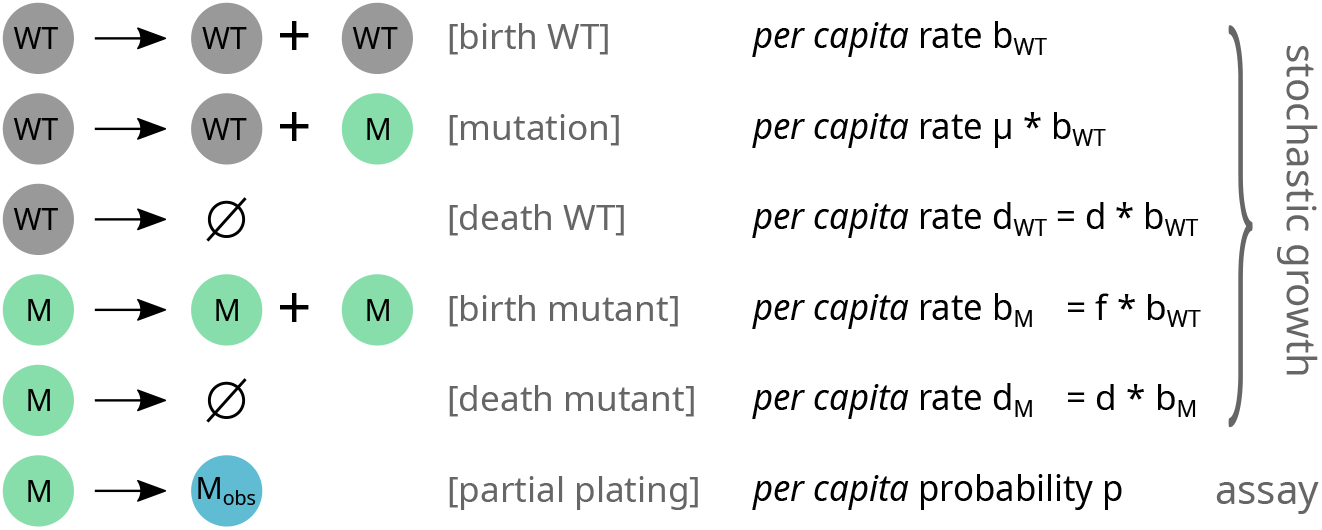
Extended fluctuation test model. *WT* and *M* represent wild-type and mutant individuals (as classical in fluctuation tests, a single class of mutants is considered). *M*_*obs*_ represents the mutants which will be observed when assaying the population by selective plating, and is different from *M* in case of partial plating (only a sample *p* of the population is plated). *µ* is the mutation rate per genome per division. Birth and death rates of the mutant and the wild-type are linked by *b*_*M*_ ≠ *b*_*WT*_ *× f* and *d*_*M*_ ≠ *d*_*WT*_ *× f* where *f* is the fitness effect of the mutation.

Other variables are the same as in the standard fluctuation test: mutation rate (*µ*) per cell division, initial and final total population sizes *N*_0_ and *N*, and observed number of mutants in the final population (output of the model). The growth dynamics can be piecewise-constant, with death and mutation rate parameters changing for each growth phase. The standard fluctuation test model can be recovered from our model by taking *p* = 1, *f* = 1, and *d* = 0, as empirically verified further below (Figure S1a).

We developed a fast simulator for this model, based on a mix of stochastic and deterministic techniques: wild-type birth and death are approximated with a deterministic model, while mutation events and mutant birth and death are simulated by a stochastic model. This simulator is further described in section 5.1.

To validate the accuracy of this simulator, we compared its outputs (number of mutants for several replicate populations) to those obtained with (1) other tools based on analytical methods in the restricted case where these methods apply, and (2) the slower Gillespie algorithm (reference stochastic simulation method) in the general case. The analytical reference methods that we used were the recursion formula proposed by Ma, Sandri and Sarkar (Ma et al., 1992), the Mandelbrot-Koch (Koch, 1982; Mandelbrot, 1974) method implemented in rSalvador, and the ‘`rlfan`’ function provided by Flan. As seen in Figures S1, S2, S3, our simulator not only yields statistically indistinguishable results from the analytical methods and the standard simulation method, but also provides a several orders of magnitude speedup over the standard simulation method (Gillespie algorithm).

Note that not all features of this forward simulator are exploited in the parameter inference methods described further below. In particular, so far the parameter inference in multi-phase growth models is restricted to two-phase growth with variable death rate but constant mutation rate.

## 3 Development of a simulation-based parameter inference method

Based on this forward simulator, we developed a likelihood-free Approximate Bayesian Computation parameter estimation method, described in section 5.2 of the Appendix. Briefly, a Metropolis-Hastings algorithm is used to walk through the parameter space, the acceptance of a new parameter set being based on the resemblance between the simulated distribution of the number of observed mutants under this parameter set and the observed experimental distribution. This resemblance is determined using the statistic of Kolmogorov-Smirnov two-samples test.

Further details on the implementation and benchmarking of this method are provided in the Appendix (section 5).

### 3.1 Mutation rate inference in standard experimental conditions

We first validated this proposed parameter inference method on the standard scenario where the mutation rate is inferred in absence of death, fitness effect of the mutation or partial / imperfect plating. Using the forward simulator, we performed 750 simulations of fluctuation assays with 24 replicate populations (*r*) for each. Mutation rate (*µ*) and total cell numbers (*N*) were randomly drawn from a broad parameter range chosen to ensure a realistic number of observed mutants (see section 5.3 for details). The initial number of cells (*N*_0_) was set to 1. Other parameters were set to their default values to match the standard assumptions (relative fitness *f* =1, death rate *d*=0, plating fraction *p*=1). The parameter *µ* was drawn from a log-uniform distribution over the range [10^*−*10^, 10^*−*6^], and *N* was subsequently drawn from a log-uniform distribution over the range [10^*−*2^, 10^2^]*/µ*.

For each simulated fluctuation test, the output number of mutants for each replicate simulation and the final total population size *N* were given to the estimator, which attempted to infer *µ* based on these data. Estimated values of *µ* were then compared with the true values used as input of the fluctuation test simulation. In addition to our proposed method, we also tested three recent, state of the art estimation methods (see section 5.4 for details): rSalvador (Zheng, 2017), which implement the Mandelbrot-Koch model (Koch, 1982; Mandelbrot, 1974) as well as the standard Lea and Coulson model (Lea & Coulson, 1949) as formulated by Ma, Sandri and Sarkar (Ma et al., 1992), and Flan (Mazoyer et al. (2017), we use the ‘generating functions’ method), that are the two most widely used software in recent literature; as well as mlemur (Łazowski, 2023), which is more recent and offers additional abilities compared to Flan and rSalvador. This allows us to compare the accuracy of our simulation-based method with analytical methods (in the restricted range of scenarios where these analytical methods work). This pipeline is represented in Figure 2. Unless otherwise specified, we always report the relative estimation error value for each parameter, defined as |(true_value - estimated_value)/true value|.

**Figure 2:**
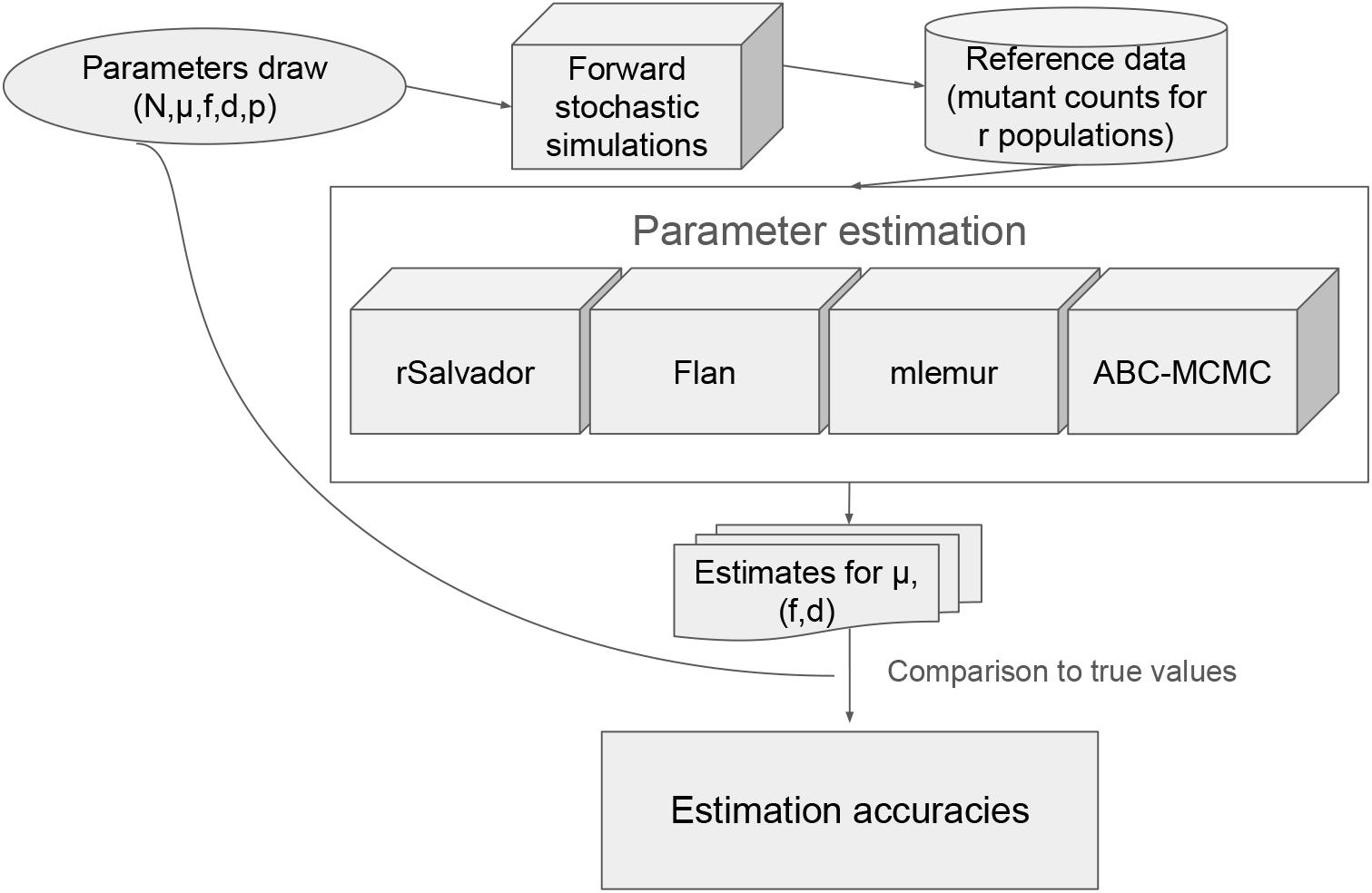
Pipeline of the accuracy test. Total cell number *N* is always provided to the estimator. Mutation rate *µ* is never provided to the estimator. Relative fitness of the mutant *f*, death rate *d* and plating efficiency *p* are sometimes provided to the estimator (as in Figures 3, 4, 5), and sometimes unknown to the estimator and jointly inferred with the mutation rate (other Figures).

As seen in Figures 3(a) and S4, in these standard conditions all methods present similar and reasonably low estimation errors, with marginal differences between tools. The mean relative estimation errors are 0.152, 0.170, 0.152, 0.169 for rSalvador, Flan, mlemur and ABC-MCMC respectively. This slight difference in accuracy between tools is small but statistically significant according to a non-parametric ANOVA (Kruskal-Wallis p-value = 0.0027, Dunn p-value for rSalvador-Flan and mlemur-Flan*<*0.01).

**Figure 3:**
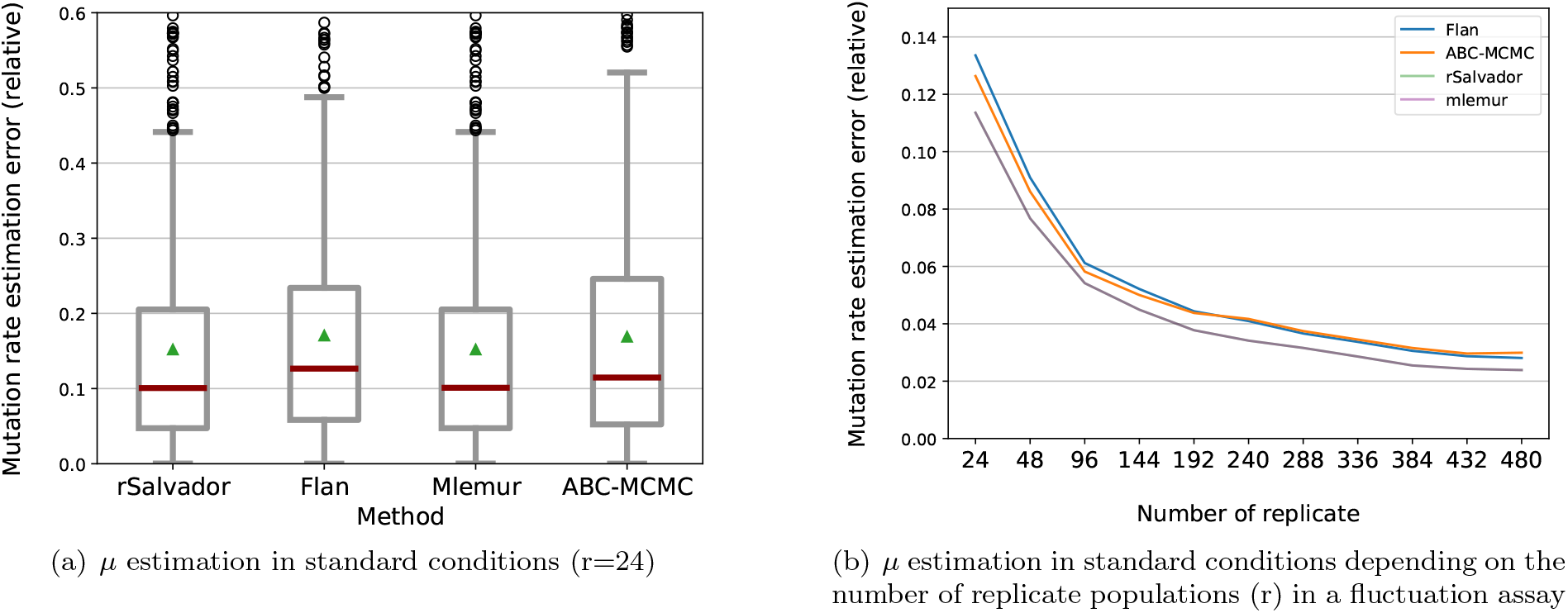
Mutation rate estimation in standard conditions. **(a)** Mutation rate (*µ*) estimation errors through rSalvador, Flan, mlemur, and ABC-MCMC (our proposed method). *µ* is the only unknown parameter, other parameters were set to match the standard assumptions: no fitness difference between mutants and wild types (*f* =1), no death (*d*=0), full population plated without sampling and every mutant detected (*p*=1). Medians are shown with red lines, means are shown with green triangles, the grey boxes represent the 25th and 75th percentile. In this simple scenario, rSalvador and mlemur implement the same method, the outputs are thus perfectly similar. **(b)** Effect of the number of replicate populations on mutation rate estimation accuracy. We simulate 486 fluctuation assay data sets with random *µ* and random N as above with 480 replicates (*r*), but only provide a sample of the replicates (varying from 24 to 480) to the estimators. For each number of replicate populations, we report the mean estimation error for the 486 datasets.

In this simple setup, we further investigated the main sources of estimation errors. We found no evidence that the true *µ* values significantly impact the accuracy of the inference (Figure S5), but we found that for all methods, the estimation error increases as the median of mutant counts decreases (Figure S6). This is easily understood as these empirical distributions with low mutant counts are not as smooth. We also observe that estimation errors are strongly correlated between all methods for a given dataset (Figure S7). These observations support the idea that in this simple setup, the accuracy is limited by the data more than the method.

To further explore how estimation accuracy is limited by the available experimental data, we varied the number of replicate populations (r) in the simulated fluctuation test experiments. As expected, for all four methods, estimation error decreases when increasing the number of replicate populations (Figure 3(b)). All tools kept showing comparable performance, with rSalvador and mlemur being marginally more accurate.

### 3.2 Mutation rate inference in known, non-standard conditions

Mutations assessed in a fluctuation assay may induce a fitness cost for the mutant bacteria, as it is for example often the case for antibiotic resistance mutations in absence of antibiotic (Andersson et al., 2007; Melnyk et al., 2015; Reynolds, 2000; Vogwill et al., 2016). When this cost is known, rSalvador, Flan and mlemur are able to take into account a known differential fitness of the mutant (relative fitness of the mutant compared to the wild-type *f* ≠ 1).

Another recurrent problem in the fluctuation assay is the plating efficiency / plating fraction. Plating may be imperfect, in the sense that not all resistant cells would be poured and survive on the selective medium, or the population may be purposely diluted before plating. Consequently, recent tools propose a correction for a known sampling factor (*p <* 1).

We thus tested our proposed estimator as well as rSalvador, Flan and mlemur in these conditions where *f* or *p* (or both) differ from 1 but have known values (see section 5.3 for draw of parameter values). As seen in panels (a) and (b) of Figure 4, we found that all four estimators correctly infer the mutation rate. Relative errors are overall similar (mean value around 0.06), the four methods presenting no significant difference when *f* ≠ 1 (Kruskal-Wallis p-value = 0.058), and Flan being marginally less accurate than every other tools when *p <* 1 (Dunn test p-value *<* 0.02).

**Figure 4:**
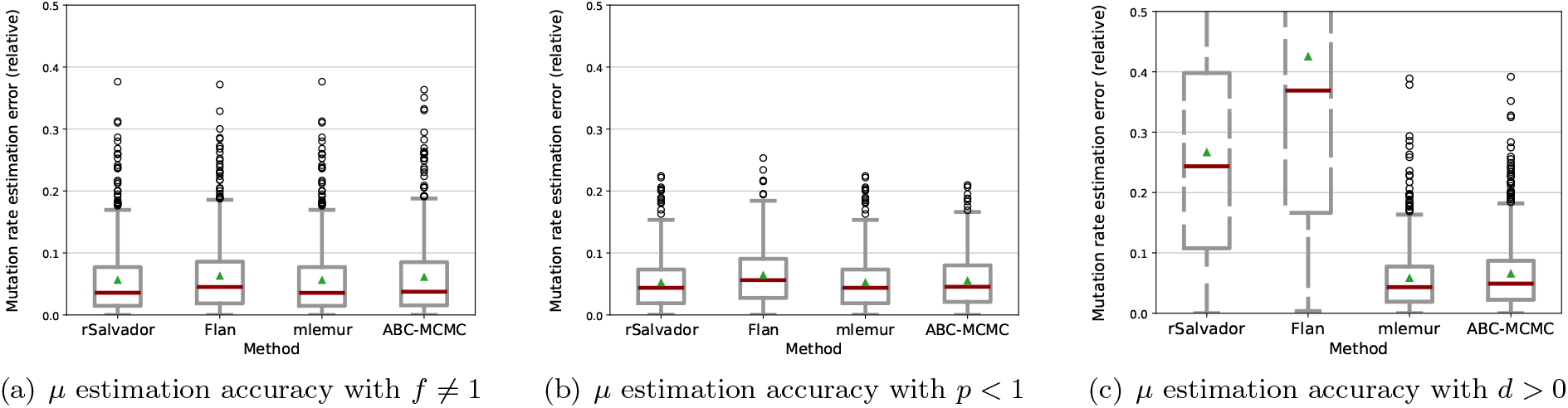
Mutation rate estimation accuracy when departing from the standard model with a single, known parameter. This corresponds to differential fitness of the mutant *f* (panel a), death rate *d* (panel b) or sampling fraction *p* (panel c). **(a)** We simulated 500 fluctuation assays with mutation rate *µ*, fitness of the mutant *f* and population size N drawn in a broad space of realistic values (see section 5.3) with *r* = 100 replicate populations. The value of *f* is provided to the estimator. Other parameters were set to match the historical, standard assumptions (no death : *d* = 0, full population plated without sampling and every mutant detected : *p* = 1). **(b)** We simulated 430 fluctuation assays, as above but with sampling fraction *p* randomly drawn and provided to the estimator (and mutant fitness back to the default *f* = 1). **(c)** We simulated 571 fluctuation assay, as above but with death rate *d* randomly drawn and provided to the estimator (and mutant fitness back to the default *f* = 1, sampling fraction back to the default *p* = 1). As rSalvador and Flan are not designed to consider this scenario, we display the results obtained with these tools in dashed gray.

An important limitation of most methods is their inability to take into account cell mortality during the experiment. Neither Flan nor rSalvador can take into account cell death (Flan has an implementation to take into account death of the mutant but not of the wild-type). Mlemur proposes a simple correction term for constant cell-death. By proposing a simulation-based approach, we aim to provide a method which can take cell mortality into account, at a non-constant rate as observed in experimental data (Frenoy & Bonhoeffer, 2018).

We first tested that our method can correctly infer the mutation rate in conditions where *d* differs from 0 but has a known, constant value. As seen in panel (c) of Figure 4, both ABC-MCMC and mlemur successfully estimate the mutation rate with accuracy remaining at the same level as previously (with r = 100), with mean relative errors of 0.065 and 0.058 respectively, mlemur being slightly more accurate (Dunn test p-value *<* 0.01). rSalvador and Flan cannot account for cell mortality. As they are nevertheless used in real, imperfect setups where hypotheses of the models are not always met, we include them in the benchmark to see how much unaccounted cell death impacts mutation rate estimation accuracy. As expected, they display a much higher estimation error in these conditions.

We then tested whether our proposed method can infer *µ* under the most complex experimental conditions where the three aforementioned difficulties are present simultaneously, that is to say when there is a fitness effect of the mutation (*f* ≠ 1), cell mortality in the culture (*d >* 0), and a fraction of the total culture is assayed (*p <* 1). Results (Figure 5(a)) show that both mlemur and ABC-MCMC succeed with a comparable relative error as before, around 0.05. As expected, rSalvador and Flan are not accurate in this scenario. This shows that ABC-MCMC and mlemur can both accurately infer *µ* in the presence of constant cell death, with a fitness effect of the mutation, and with sampling before plating, as long as the parameters describing death, fitness and sampling are known and taken into account.

**Figure 5:**
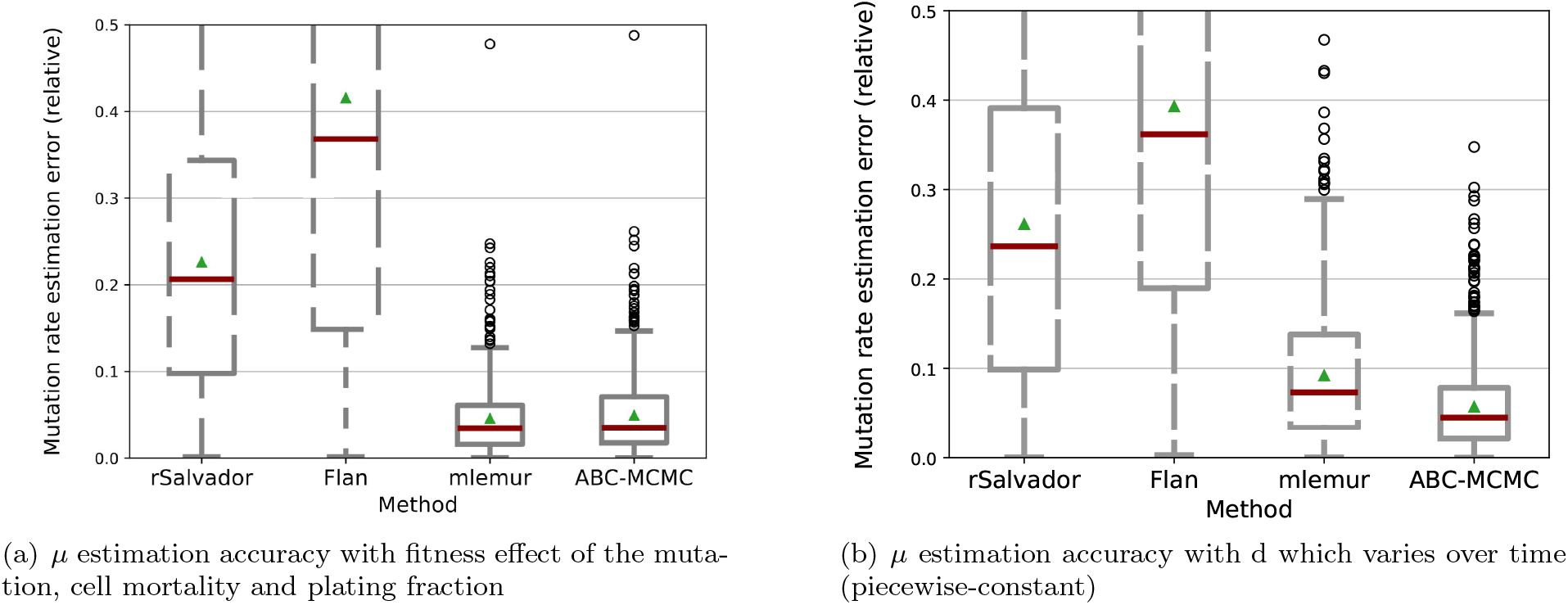
Mutation rate estimation accuracy when departing from the standard model with several known parameters. **(a)** Three demographic parameters (fitness effect *f*, death rate *d*, and plating fraction *p*) have a non-default but known value. We simulated 750 fluctuation assays with *r* = 100 replicate populations. **(b)** Death rate changes over time (two growth phases with different, known *d* values). We simulated 1000 fluctuation assays with *r* = 100 replicate populations. Two *d* values were drawn, corresponding to two distinct phases with different cell mortality. We defined two *N* values, *N*_*i*_ and *N*_*f*_, respectively corresponding to the number of cells when the population enters the second growth phase, and the final number of cells after the end of the second growth phase. *N*_*i*_ and *N*_*f*_ were respectively set to 10^5^ and 10^8^.

We finally tested the situation where cell mortality is not constant and varies during the growth phase. This matches experimental observations showing multiphase population dynamics with non-constant cell death under stress (Frenoy & Bonhoeffer, 2018; Grant et al., 2008; Hart et al., 2019), but also specific fluctuation assay protocols where cells are first grown under stress, then recover in a stress-free medium, before selective plating (López et al., 2007). In this context, we generated a dataset with two growth phases, having the same *µ* but different *d* values. We then inferred *µ*, given the different *d* values and the population size at which this value changed. The results are shown in Figure 5(b). Here again ABC-MCMC correctly estimates the mutation rate, with a mean relative error comparable to previous results (0.056). rSalvador and Flan are highly inaccurate as expected as they do not consider cell death. mlemur can consider constant cell death but not piecewise-constant, and was given the average cell death parameter between growth phase. It displayed a higher accuracy than rSalvador and flan, but lower than ABC-MCMC. We furthermore tested the same setup (inference of *µ* with known, piecewise-constant *d*), with the addition of a known fitness effect of the mutation and a known sampling factor. ABC-MCMC still succeeds, with similar precision (Figure S8).

### 3.3 Two parameters inference

Theoretically speaking, ABC-MCMC methods can simultaneously infer an arbitrarily high number of parameters, under two conditions: (1) identifiability – all parameters need to have significant and non-overlapping effects on the mutant count distribution, and (2) sufficiently large amount of data – number of replicate populations in our case. We tested simultaneous inference of *µ* and other parameters (relative fitness of the mutant, and death rate), and report the results in the following section.

#### 3.3.1 Mutation rate and Relative fitness

Mutation rate and relative fitness joint inference have already been implemented in analytical tools such as rSalvador and Flan, which was an important innovation compared to previous tools such as Falcor (Hall et al., 2009) implementing the MSS-LME. Here we tested ABC-MCMC joint inference of these two parameters, and compare it with results obtained with rSalvador and Flan. mlemur was excluded from this analysis as it cannot infer the relative fitness of the mutant.

We evaluated the three methods in the scenario where *f* ≠ 1 and is unknown. We performed 1943 simulations of fluctuation assays, with *r* = 100 replicate populations for each. *µ, N*, and *f* were randomly drawn over parameter ranges which ensure a realistic number of observed mutants (see section 5.3 for details). *µ* and *N* were drawn over the same range as before, *f* was drawn uniformly over the range [0.4, 1.2]. Other parameters were set to their default values to match the standard assumptions (death rate *d* = 0, plating fraction *p* = 1). For each simulated fluctuation test, the output number of mutants for each replicate simulation and *N* were given to the estimator, which attempted to simultaneously infer *µ* and *f* based on these data.

Results are shown in Figures 6 and S9. In these conditions, we observe roughly similar estimation accuracies for the three methods. *µ* mean relative estimation errors (panel (a)) are 0.072, 0.077 and 0.092 for rSalvador, Flan and ABC-MCMC respectively. *f* mean relative estimation errors (panel(b)) are 0.062, 0.072 and 0.068 for rSalvador, Flan and ABC-MCMC respectively. rSalvador and Flan are marginally more accurate for estimating *µ* compared to ABC-MCMC (p-value *<* 0.01). For *f* inference, rSalvador is marginally more accurate than the two other tools (p-value *<* 0.05). Moreover, rSalvador estimation errors have a slightly lower variance. ABC-MCMC displays a slightly higher variance for *µ* estimation errors, and Flan a slightly higher variance for *f* estimation errors.

**Figure 6:**
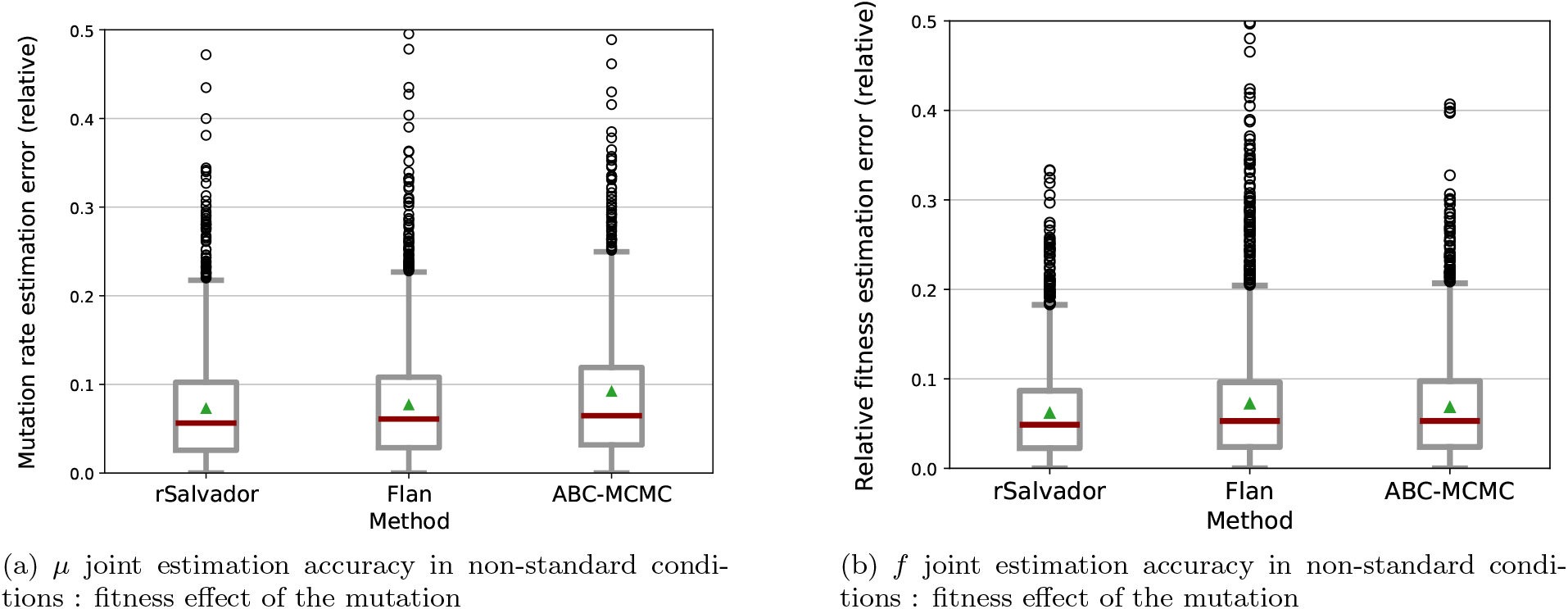
Joint *µ* and *f* estimation, for rSalvador, Flan and ABC-MCMC. Panel (a) shows *µ* estimation errors; panel (b) shows *f* estimation errors. We simulated 1943 fluctuation assay data sets with mutation rate *µ, f* and N randomly drawn, *r* = 100 replicate populations, and jointly inferred *µ* and *f*. Other parameters were set to match the historical, standard assumptions (death rate *d* = 0, sampling fraction *p* = 1).

ABC-MCMC displays a few rare but strong high outliers values (better shown in Figure S10(a)), as expected from a stochastic method, hence the higher average of *µ* estimation error while the median is similar than with the other tools. rSalvador often fails to converge with default parameters (Figure S10(b)), but most (although not all) convergence failures are easily solvable by providing a different starting point value for the parameter search.

Mean estimation errors are, here again, higher on data sets with small mutant counts (in particular when the median number of mutants is lower than four) for the inference of both parameters (Figure S11). The estimation accuracy of *µ* is not significantly impacted by the true *µ* value (Figure S12). This result is consistent with what we previously observed.

Joint distributions of the errors for *µ* and *f* are shown in Figure S13, without the absolute value ((true value - estimated value)/true value). *µ* and *f* estimation errors are negatively correlated with all methods, as expected (an underestimation of the mutation rate is compensated by an overestimation of the fitness of the mutant, and vice versa). These correlations are stronger for rSalvador and Flan than for ABC-MCMC, but are always statistically significant.

Adding a second parameter to infer along with *µ* slightly decreases the estimation accuracy as intuitively expected, but the estimation remains sufficiently accurate with *r* = 100 replicate populations. We show in Figure S14 the effect of this number of replicate population *r* on the accuracy of joint *µ* and *f* estimation. We find that running *r* = 24 replicate experimental populations as classically done gives a considerably higher estimation error than when inferring a single parameter.

We further characterized how each of the methods behave depending on true *f* value (Figure S15). Focusing on *µ* inference (panel (a)), we can see a clear proportional relationship between *f* value and *µ* estimation errors: *µ* estimation error increases with *f*, the slope of this relationship being slightly higher for ABC-MCMC. For *f* estimation (panel (b)), the error overall stays stable as *f* increases, although ABC-MCMC seems to display a small increase in accuracy as *f* increases (Figure S16). These results overall show that as mutant fitness increases, it is increasingly difficult for all estimators to infer *µ*, however this loss of accuracy is not correlated with a worst *f* inference.

Finally, we repeated the latest test with datasets which were simulated with *f* =1 (no fitness effect), to ensure that the tools would not falsely infer a non-existing fitness effect (Figure S17). With 880 simulations, we observe similar results than with previous conditions: the three methods display close estimation errors, with reasonable values (between 0.082 and 0.097 for *µ* and between 0.064 and 0.080 for *f*) with a small advantage for rSalvador (Dunn test p-value *<* 0.01). The same observations on variability are also reported. This shows that the two parameters inference versions of Flan, rSalvador and ABC-MCMC are safe to use even in situations where *f* = 1.

Finally, we used ABC-MCMC to test *µ* and *f* joint inference in the case where there was cell mortality (with a known, constant death rate *d* ≠ 0, Figure S18) and in the case where there was additionally sampling before plating (known sampling fraction *p* ≠ 1 and known death rate *d* ≠ 0, Figure 7). Both estimates of *µ* and *f* present comparable results to previous sections (mean estimation error 0.08 for both). These results show that ABC-MCMC is able to precisely infer *µ* and *f* together while taking cell death and partial plating into account, as long as *d* and *p* are known. Estimations using rSalvador and Flan were also included for comparison, following the same rationale as before: they are not designed to work in these conditions, but are nevertheless used in imperfect conditions where all hypotheses are not always met. As expected, these methods fail to infer *µ*, with a very large error. Surprisingly, Flan is nevertheless able to correctly infer *f*.

**Figure 7:**
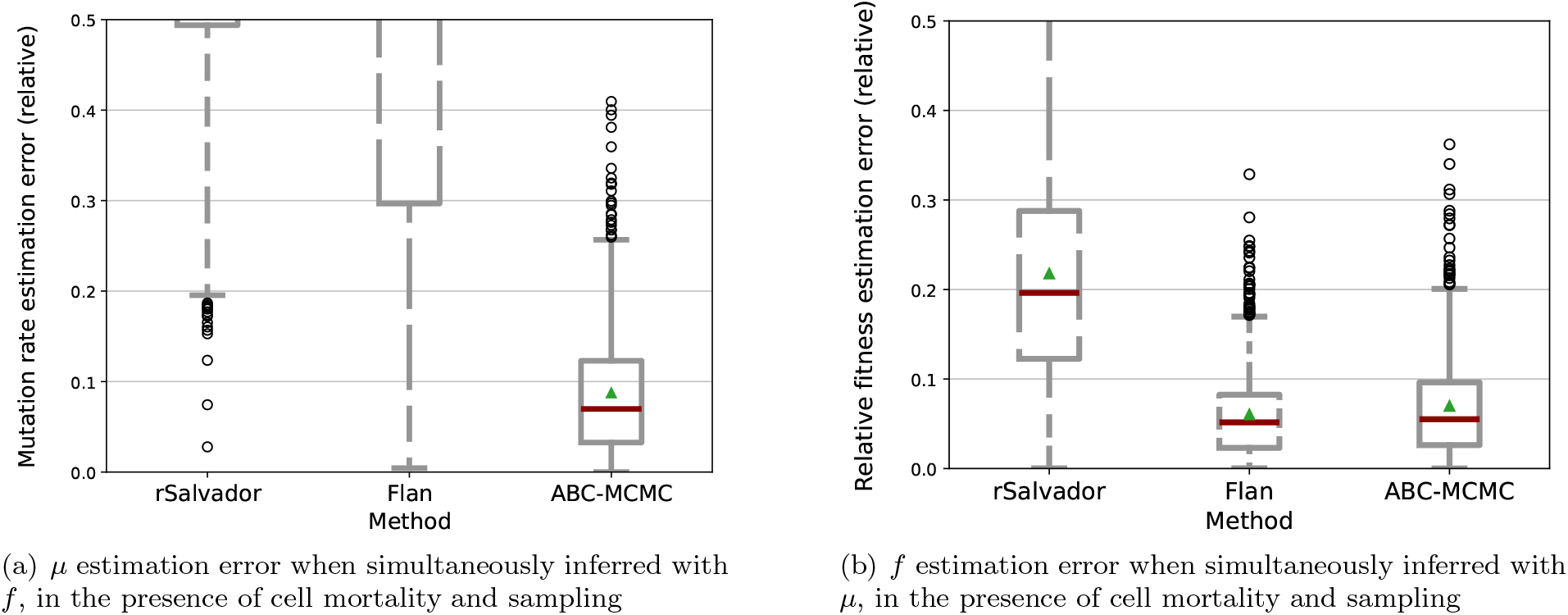
Joint *µ* and *f* inference in non-standard conditions: known cell mortality (*d* ≠ 0) and sampling (*p* ≠ 1). We simulated 915 fluctuation assay data sets with *µ, f, d, p* and *N* randomly drawn and *r* = 100 replicate populations, and jointly inferred *µ* and *f* with ABC-MCMC, giving the known *d* and *p* values to the estimator. rSalvador and Flan were included for comparison purposes. rSalvador does not have a function to take into account both *f* and *p*, hence, we jointly inferred *µ* and *f* with *p* value assumed to be 1.

In conclusion, *µ* and *f* can reliably be jointly inferred by Flan, rSalvador and ABC-MCMC in simple conditions (no death and sampling). In more complex conditions (known death and sampling fraction), it can be reliably inferred by ABC-MCMC, and not by any of the other existing tools.

#### 3.3.2 Mutation rate and Death rate

We then tested ABC-MCMC’s ability to infer *µ* and *d* simultaneously, with a constant *d* ≠ 0. We performed 370 simulations of fluctuation assays, with *r* = 100 replicate populations for each, and randomly drawn *µ, d* and *N* (see section 5.3 for details). Other parameters were set to their default values to match the standard assumptions (see Box 1: relative fitness *f* = 1, plating fraction *p* = 1). For each simulated fluctuation test, the output number of mutants for each replicate simulation and *N* were given to the estimator, which attempted to simultaneously infer *µ* and *d* based on these data. As shown in Figures S19, S20, *µ* estimation was highly unreliable, with a mean relative error 0.34 and a high variance between replicates. The same observations are made for *d* estimation, with a mean relative error 2.0 driven by strong outliers.

Increasing the number of replicate experimental populations, increasing the number of simulations at each MCMC step or increasing the length of the MCMC itself did not provide better results. We tried up to 500 replicate experimental populations in the reference dataset, 3000 simulated populations at each step, and 10000 steps in the MCMC, without improving the results.

To understand why this joint inference of the mutation rate and the death rate fails, we plotted the landscape of distances between a reference simulation and the simulations obtained for all combinations of *µ* and *d* (Figure S21a). We found that many different parameter pairs equally fit the reference distribution, showing poor identifiability of *µ* and *d*. As a comparison, the same landscape is shown in Figure S21b for *µ* and *f*, for which the inference succeeds, and shows as expected a clear, well-delimited zone of low Kolmogorov-Smirnov (KS) distances. Said otherwise, cell mortality has an impact on the mutant count distribution that can not be disentangled from the effect of the mutation rate, unlike the one from the fitness of the mutant.

## 4 Discussion

Classical methods used for mutation rate computation are based on mathematical models. These have the advantage of being very fast and precise, as long as the experimental setup respects a certain number of necessary conditions assumed by the model. However, it is increasingly clear that this list of hypotheses does not always reflect the reality of bacterial growth. In this paper, we present a new simulation-based approach for mutation rate estimation in variants of Luria and Delbrück’s fluctuation assay. This method allows taking into account or estimating a large number of secondary parameters that shape the final mutant count distribution of a fluctuation assay. So far, these parameters include fitness effect of the mutation, sampling before selective assay (partial or imperfect plating), and mortality (with a piecewise-constant death rate). The backbone of our method is a fast stochastic, forward in time simulator of the experiment. Coupled with an ABC-MCMC method based on the Metropolis-Hastings algorithm, we showed that we were able to infer the mutation rate (as well as the fitness of the mutant when relevant) in a large set of experimental conditions. All the parameter inference scenarios tested in this article are summarized in Table 1, with an indication of which tool provides reliable results for each of these scenarios.

**Table 1:** Summary of conditions where our proposed method and existing tools can be used.

| Condition |  |  |  | Tool |  |  |  |  |
| --- | --- | --- | --- | --- | --- | --- | --- | --- |
| mrate | f | d | p | ABC-MCMC<br>(proposed) | rSalvador | Flan | mlemur | Figure |
| Infer | 1 | 0 | 1 | OK | OK | OK | OK | 3(a) |
| Infer | Known ≠ 1 | 0 | 1 | OK | OK | OK | OK | 4(a) |
| Infer | 1 | Known > 0 | 1 | OK | X | X | OK | 4(b) |
| Infer | 1 | 0 | Known < 1 | OK | OK | OK | OK | 4(c) |
| Infer | Known ≠ 1 | Known > 0 | Known < 1 | OK | X | X | OK | 5(a) |
| Infer | 1 | Known piecewise cte | 1 | OK | X | X | X | 5(b) |
| Infer | Known ≠ 1 | Known piecewise cte | Known < 1 | OK | X | X | X | S8 |
| Infer | Infer | 0 | 1 | OK | OK | OK | X | 6 |
| Infer | Infer | Known > 0 | 1 | OK | X | X | X | S18 |
| Infer | Infer | Known > 0 | Known < 1 | OK | X | X | X | 7 |
| Infer | 1 | Infer | 1 | X | X | X | X | S19 |

## 4.1 Comparing parameter estimation accuracy between tools

The four tested methods yield comparable results in simple conditions (estimation of *µ* in absence of fitness effect, death or sampling), with rSalvador and mlemur having a marginally better accuracy than Flan and ABC-MCMC.

In addition to producing similar estimation errors in average, we note that the four tools tend to produce the same error for a given dataset. This shows that the accuracy of the estimation is mostly limited by the experimental data. More specifically, two properties directly impact estimation accuracy of all four methods: the number of replicate experimental populations (*r*), and the median number of observed mutants. For all methods, when the median of mutant count is low (*<* 4) there is a significant decrease in accuracy. This is due to the distributions containing many zeros and small values, which limits the ability to identify mutation rate. Additionally, as expected, we observed a significant improvement in estimation accuracy when the number of replicates increases. In particular, the gain is significant when increasing the number of replicates from 24 (classically performed in the lab) to 192, with a more marginal gain when yet more replicates are provided. Compared to 24 populations (classically used in experimental data), estimation errors were approximately half as large for 96 replicates and three times lower for 192 replicates. This observation was consistent for the four tested methods.

In the simple conditions shown in Figure 3 (estimation of *µ* in absence of fitness effect, death or sampling), simulation-based methods have no particular added value, and these conditions were rather used as a first check of our proposed method. It is expected to see an analytical method like rSalvador, Flan or mlemur being more accurate than a simulation-based method like ABC-MCMC because of the stochasticity aspects involved in ABC processes. Nevertheless, this difference in accuracy is relatively small, making ABC-MCMC a possible choice when inferring the mutation rate under these conditions where analytical tools are favored due to their speed.

The main advantage of ABC-MCMC is the capacity to be used in a broader range of specific biological scenarios that analytical methods cannot always consider, as discussed in the next paragraphs.

### 4.2 Extensions of the historical Luria & Delbrück model: considered biological scenarios

We tested ABC-MCMC and the three reference analytical tools in a large set of situations which have historically needed further mathematical development beyond the original growth model from Luria and Delbrück, as well as in some situations where current analytical methods fail.

The first complication to consider is the fitness effect of the mutation. While the original model from Lea and Coulson assumed equal fitness of the wild-type and the mutant, fluctuation tests are often performed on antibiotic resistance mutations which are classically associated with a fitness cost in absence of antibiotic (Andersson et al., 2007; Levin et al., 1997; Melnyk et al., 2015; Vogwill et al., 2016). This scenario has been considered in a large body of literature and prompted further mathematical development, such as Mandelbrot-Koch model (Koch, 1982; Mandelbrot, 1974), which allows the fitness effect of the mutation to be taken account or to be jointly inferred with the mutation rate. All four tools tested in this work are able to take into account fitness effect (positive or negative) of the mutation when estimating the mutation rate. Flan, rSalvador and ABC-MCMC can infer the fitness effect when unknown, with accuracies similar to mutation rate, while mlemur cannot.

Another complication is cell mortality, which was historically less well studied and often neglected by mathematical models. Death rates are considered negligible for growth in rich medium, but mortality is likely to occur at a high rate when the assay is performed under stressful conditions, such as low doses of antibiotics (Frenoy & Bonhoeffer, 2018). Mathematically accounting for death has proven to be more complex than for fitness effect. Flan implements a model where only mutant cells die, which is not suitable for the generally considered scenario. Mlemur suggests a correction term which only works when death rate is constant throughout the experiment, which does not match empirical evidence. Experimental data quantifying microbial death are scarce, but show that death rates are not constant (for example for *Escherichia coli* under antibiotic stress (Frenoy & Bonhoeffer, 2018), *Saccharomyces cerevisiae* under lysine depletion (Hart et al., 2019), or *Salmonella enterica* within mice hosts (Grant et al., 2008)), hence our choice of considering a piecewise-constant death rate. Mlemur can infer the mutation rate when cell death is constant and the fitness effect of the mutation is known. rSalvador cannot take mortality into account at all, and Flan only considers mortality of mutant cells but not of wild-type cells, these two tools are thus not suitable for this scenario. We show that ABC-MCMC can accurately infer the mutation rate in this scenario characterized by a more complex (two-phase) cell demographics as well as in the presence of unknown fitness effect of the mutation. But contrary to fitness effect, we found that it can not jointly infer an unknown death rate with the mutation rate: Figures S19, S20 and S21 show that these two parameters are not identifiable (at least for the statistical measures we consider). That known, to improve *µ* inference in these conditions, fluctuation assays should be coupled with other experiments to determine cell mortality (Frenoy & Bonhoeffer, 2018; Golchin et al., 2012; Mahmutovic et al., 2021; Myhrvold et al., 2015).

The final departure from the original model that we considered is sampling before selective plating (or partial plating). This sampling can be implemented by the experimenter to lower mutant count to numbers that can be counted on a Petri dish, when mutation rates are high or the mutational target is large. But it can also result from what was historically referred to as imperfect plating (the plating procedure may lose some cells or cause them to die before they can form a colony). Mathematically speaking, both mechanisms lead to the same model. Taking into account a known sampling factor has historically proven surprisingly complex, with an inaccurate correction term (for example proposed by Falcor, Hall et al. (2009)) being used in a large set of papers (as discussed by Zheng (2017)). The four recent tools tested in this article are able to infer the mutation rate in the presence of a known sampling factor, with similar accuracies. ABC-MCMC, mlemur and Flan can do so in the presence of a known fitness effect, while rSalvador can not.

As summarized in Table 1, our proposed method successfully infers mutation rates in the presence of all these factors extending the original Luria & Delbrück model (potentially unknown fitness effect of the mutation, sampling before selective assay, piecewise-constant death rate), and is to our knowledge the only method able to do so.

As further discussed below, our simulation-based method could in principle accommodate other departures from the original model stemming from recently emerged hypotheses concerning the molecular aspects of mutagenesis, beyond the scope of the present work.

### 4.3 Potential limitations of the proposed method

The main advantage of ABC-MCMC is its ability to consider complex growth scenario where other methods fail, due to its simulation-based nature. But as a result of this simulation-based nature, it was anticipated that ABC-MCMC estimation errors would be higher than analytical methods by nature for simple growth conditions where analytical methods can be used. Although these differences exist, they are small in our benchmark.

Beyond accuracy, this method is nonetheless computationally more costly as it relies on stochastic simulations. Important factors determining computation time include the number of steps used in the chain of the Metropolis-Hastings, the number of replicate populations simulated at each step, and parameter values for those simulations. In particular, simulating populations with large mutant counts takes more time, as the mutants demographic is computed using a “standard” stochastic simulation scheme (Gillespie) without deterministic approximations. Overall, the runtime for the full parameter estimation process for one experimental dataset can vary from a couple of seconds in most cases to several minutes in less favorable cases (Figure S22). These unfavorable cases were for example encountered for simulations with large *f* values, which lead to a large number of mutants, making the simulator extremely slow. It should however be noted that experimental datasets are characterized by a number of mutants which can be counted on a Petri dish (no more than a thousand). Parameter draw in the MCMC leading to such large mutant counts are thus going to be eventually discarded as they are very far from the experimental data. We thus implemented a cutoff on simulation time and/or maximal number of mutants in the simulator: when this cutoff is exceeded, the simulation is killed and the parameter draw is discarded. This simple stop condition is likely to be sufficient to avoid long simulation times without loss of accuracy for realistic datasets.

### 4.4 Parameter choices

Biological parameters for the simulated fluctuation assays (population size, mutation rate, fitness effect of the mutation, death rate) were chosen over a large range that covers values which we considered realistic based on the literature (as described in section 5.3). Because fluctuation assays score a set of mutations which confer a selectable phenotype, and not directly a specific genotypic mutation, mutation rate values can vary over several orders of magnitudes depending on the target size (number of different mutations which can confer the phenotype of interest). Death rate values are rarely estimated due to the experimental complexity to do so. Fitness effects can in principle vary over a broad range, but we may argue that a marker with a very strong fitness effect is a poor choice for a fluctuation test assay, justifying the restricted range we chose to study.

Beyond biological parameters, our MCMC method involves several computational parameters, such as the length of the chain, the burn-in period, the acceptance threshold, the transition parameters, etc. This method is thus computationally more complex than rSalvador and Flan, with more parameters to be understood in order to tune it. The default choices are suitable for most situations and were used for all results presented in this article. An expert user may nonetheless want to tune them depending on the specificity of their data, their need for accuracy in balance with computational power, etc.

### 4.5 Future directions, possible extensions of the method

We believe that a significant advantage of our method resides in its conceptual simplicity: despite being computationally expensive, the process of simulating bacterial growth and mutagenesis with a certain set of parameters and comparing the results to experimental data is conceptually very simple, and understandable without advanced mathematical knowledge. The forward simulator is easy to adapt to a large variety of biological scenarios beyond those directly considered in the article.

In the Box 1, we highlight “classical” hypotheses already identified by Foster (2006) that we identified as potentially circumventable with modified versions of our simulator.

More recent hypotheses which emerged in recent years also challenge the original model. We can for example think of mutation rate heterogeneity (Alexander et al., 2016; Engelhardt & Shakhnovich, 2019; Lansch-Justen et al., 2024; Matic, 2019; Uphoff et al., 2016), which posits that different subpopulations have a different mutation rate despite having a similar genetic background, shaped for example by stochasticity of the expression and action of repair mechanism. Another recently considered phenomenon is the phenotypic delay between the mutation and expression of the phenotype (Carballo-Pacheco et al., 2020; Sun et al., 2018), potentially influencing the mutant count distribution. We can also mention density-dependent mutation rates (Krašovec et al., 2018), or more generally the effect of stationary phase and nutritional stress (Kivisaar, 2003; Maharjan & Ferenci, 2017; Saint-Ruf et al., 2007). All these hypotheses further increase the complexity of the underlying model of population growth and mutagenesis, and potentially add more parameters to take into account when computing the mutation rate or to estimate jointly with the mutation rate. Analytical methods are limited in their ability to do so, but simulations are generally a powerful tool to consider scenario which are mathematically untractable. Simulation-based methods can be extended to account for arbitrary complications such as those discussed above as long as they can be integrated in the simulated model.

Simulation-based studies can be limited by the computational cost of the simulator. This computational cost largely arise from the discrete and stochastic aspects. Stochasticity is nevertheless an important feature, as it is at the heart of the evolutionary process, not only for mutagenesis, but also for the demographics of the mutant population which impacts mutant establishment (Alexander, 2023). The simulator that we propose treats the mutant population in a fully discrete and stochastic way, but makes more continuous and deterministic approximations for the wild-type population.

The family of simulation models that we use features a finite number of distinct populations. To go back to the example of phenotypic heterogeneity (a broad example of a complication of the mathematical models with a major evolutionary relevance), this means that it can feature a small number of subpopulations with different growth and mutagenesis properties (such as in the model proposed by Lansch-Justen et al. (2024)). But different types of simulators –based on slower algorithms, likely individual-based– would be needed to consider the effect of a more continuous heterogeneity, for example for growth and death rates which are impacted by asymmetric division (Stewart et al., 2005).

Beyond mutation rate estimation, forward simulators can be used to better understand the effect of arbitrary biological phenomena on the dynamics of emergence of mutants in bacterial populations (as for example done by Vasse et al. (2022) for antibiotic stress with a preliminary version of this simulator). These methods could also be adapted to the study of bacterial growth and mutant emergence in the human body, or to the evolution of cancer tissues, which features similar processes where the invasion of cancer cells is driven by the interplay between mutagenesis and population dynamics with complex effects of the environment.

## Data availability

Code and data used in this article will be available on Zenodo, and are available as a supplementary file during peer review to respect the double-anonymized procedure from the journal.

## Author contributions

Conceptualization: AF; Investigation: AT; Methodology: AT, AF; Writing: AT, AF.

## Acknowledgments

We express our gratitude to Arnaud Gutierrez for fruitful discussions, to Ajay Rathikumar and Raphael Malak for exploratory work conducted during their master internships, to Phuong Thao Chu for her feedback on a draft of this manuscript, to Helen Alexander and one anonymous reviewer for their constructive and detailed reviews, to the High Performance Computing center from the University of Grenoble (GRICAD) for providing the computational power needed for this study, and to the Evolution English Language Support program.

## 5 Appendix: details of the methods

### 5.1 Forward model and simulator

As depicted in Figure 1, we consider a generalized version of the fluctuation test, where a wild-type and a mutant population grow with (a) differential fitness of the wild-type and the mutant, (b) death (of both the wild-type and the mutant), and (c) sampling (only a fraction of the final population is assessed).

The simulations are initialized with a small value for the number of wild-type individuals *N*_*wt*_ and no mutant (*N*_*m*_ = 0), and continue until the total number of bacteria *N*_*wt*_ + *N*_*m*_ reaches a user-provided value *N*. This corresponds to the experimental conditions of the fluctuation test, where populations are inoculated with a small number of wild-type bacteria and grown until reaching carrying capacity. The output of the simulation is the number of mutants observed in the final population (or a sample of the final population if *p* ≠ 1). As these simulations are stochastic, when running several replicate simulations, a distribution of observed number of mutants is obtained.

As a reference method, we implemented the standard Gillespie algorithm (Gillespie, 1977) for this problem. With realistic parameters for fluctuation tests in *Escherichia coli* using antibiotic resistance markers (*N* = 5 *×* 10^9^, *µ* = 5 *×* 10^*−*10^), the simulation takes 1-3 minutes per replicate simulation on a desktop computer, which is acceptable for a forward simulation study but is unreasonably slow for simulation-based parameter inference.

We were able to speed up this process by several orders of magnitude by modifying the classical Gillespie simulation scheme, without noticeable changes in the obtained distribution (Figures S1, S2, S3). The key ingredient of our proposed simulation scheme is implementing a deterministic demographics (growth and death) for the wild-type population but a stochastic mutagenesis process and a stochastic demographics for the mutant population.

The main loop simulates each mutation event:

1. The number of divisions *Nbirth*_*wt*_ of the wild-type until the next mutation event is drawn from an exponential distribution with parameter *µ* (which is a fast approximation for a geometric distribution, accurate since *µ ≪* 1) and is rounded up to the nearest integer.
2. The number of death events for the wild-type which occurred during this waiting time to the next mutation is deterministically computed as *Ndeath*_*wt*_ = *Nbirth*_*wt ×*_ *d*_*wt*_, rounded up to the nearest integer.
3. If these numbers of death and birth events result in a total population larger than the specified final population size, then the simulation has to be stopped before the mutation. The number of births and deaths of the wild-type is deterministically adjusted to reach exactly final population size, and the numbers of births and deaths of the mutant population is then determined as below. This implies that the simulation will not exactly stop at the target final population size, but this discrepancy is minor since the mutant population is considerably smaller than the wild-type population.
4. The demographics (number of births and deaths) of the mutant population during this waiting time is computed using a standard Gillespie algorithm. The total time to simulate (expressed in number of synchronous generations) is the time needed for the wild-type population to achieve the already determined number of death and birth events, and is computed as 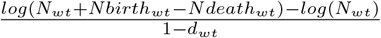 Following the standard Gillespie algorithm, the loop of this inner simulation consists in the following steps:
  - Determining the propensity of occurrence of each reaction (birth of the mutant and death of the mutant) – based on its rate per capita and on the size of the reactant population – and summing them to obtain the total reaction propensity
  - Drawing time to the next event from an exponential distribution whose parameter is the total reaction propensity, and updating the elapsed time variable. If this elapsed time becomes higher than the total time to simulate, the inner simulation is stopped.
  - Determining which of the two possible reactions occurs (random draw with probability proportional to the reaction propensity)
  - Updating the variable *N*_*m*_. In case of extinction of the mutant population, the inner simulation is stopped even if the total time to simulate has not been reached.
5. The mutation is performed (except if we determined at step 3 that the simulation has to end before the mutation) and the variables are updated.

A key underlying assumption of this algorithm is that mutations happen per cell division, and not per unit of time. The parameter *µ* is thus a probability of mutation per wild type (WT) cell division. The WT generation time is effectively used as the time unit, and all other mentioned rates (*d*_*WT*_, *b*_*M*_, *d*_*M*_) are relative to the WT generation time. This is why the “waiting time” to the next mutation event step 1 of the algorithm is also expressed in number of WT cell divisions.

This fast forward simulator is the basis of our simulation-based, likelihood-free inference method described further below. It is implemented in C / C++, and we make its source code publicly available (see section Data availability).

Several standard simplifications are made in this model, not because they are necessary for efficiency but because they are biologically relevant. The per capita rate of the WT birth reaction, formally equal to (1*− µ*) *∗ b*_*WT*_, is approximated by *b*_*WT*_, as *µ ≪* 1. The reversal mutation is always neglected, as fluctuation tests only consider the case where the mutant population is small compared to the wild-type population.

In the presence of both cell death and fitness effect of the mutation, we chose to have the same fitness effect term muliplicatively affect birth and death rates of the mutant. Our rationale behind such coupling of birth and death rates stems from experimental observations that these rates are often strongly linked in bacteria, whether between different strains (Yang et al., 2019) or between different growth conditions for the same strain (Biselli et al., 2020), as well as when considering antibiotic-induced death in different physiological conditions (Lee et al., 2018; Tuomanen et al., 1986). But we also explored an alternative model in which the mutation independently affects birth rate and death rate, and present results from this model in Figure S23.

### 5.2 Metropolis-Hastings algorithm

The proposed method for mutation rate estimation relies on the Metropolis-Hastings algorithm (Chib & Greenberg, 1995; Hastings, 1970; Robert, 2016). This algorithm uses an MCMC to look for parameter values of the simulator which produce distributions close to the experimental dataset.

The Metropolis-Hastings acceptance probability at each step of the Markov chain was determined using the Kolmogorov-Smirnov (KS) distance (Massey, 1951) between the experimental mutant count distribution and the distribution obtained by simulation with the proposed parameters at that time. This distance is a non-parametric way of comparing two distributions using their full shapes, instead of relying on summary statistics as often done in the ABC rejection scheme (Beaumont (2019) and Csilléry et al. (2010), but see Sousa et al. (2009) for an example of ABC using the full distribution instead of summary statistics).

Acceptance probability was more precisely computed as the probability that the quotient of the KS distance between the actual value and the next proposed value was inferior to a random integer uniformly drawn in the range [1, *acceptance threshold*], where *acceptance threshold* is a parameter which tunes how “exploratory” is the algorithm. default value was set as 3.

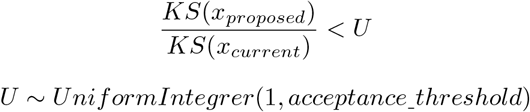

The new proposed parameters at each step of the Markov chain were randomly drawn over a normal (Gaussian) distribution, centered on the current parameter value, with a width (standard deviation) depending on each parameter. For *µ*, the width of the distribution was set as the center (mean) value multiplied by 5, for *f* and *d*, the width was set as 0.15 and 0.10 respectively.

The prior distribution for each parameter to be drawn in was determined based on the realistic range reported in the literature. The mutation rate (*µ*) range was set within [10^*−*11^, 10^*−*6^], *f* values were set within [0.4, 1.2] and *d* values were set within [0, 0.9]. These ranges are parameters which can be adjusted in the program.

This algorithm may occasionally call the simulator with unrealistic parameter values that produce a very large number of mutants, and subsequently reject the proposal as it poorly fits the reference data. This situation does not affect the accuracy, but significantly increases the running time of the algorithm, as simulations with a large count of mutants are slower. We thus implemented two simple cutoffs. First, a simulation within the MCMC is killed if it exceeds a certain duration (set to 30 seconds by default). Second, if the 3rd quartile of the mutant count exceeded 1500 in an MCMC proposal, it was automatically rejected, as such large number of mutants can not be counted on a Petri Dish.

### 5.3 Parameter draw and production of ‘experimental’ datasets

For each simulation of a fluctuation test, *µ* was first drawn from a log-uniform distribution over the range [10^*−*10^, 10^*−*6^]. *N* was subsequently drawn from a log-uniform distribution over the range [10^*−*2^, 10^2^]*/µ*. Drawing N over a range which depends on *µ* is necessary to obtain a realistic number of mutants. In a real experiment, this number will be counted on a Petri dish, and counts higher than*≈* 500 *−* 1000 will be discarded as it will not be possible to count individual colonies. Conversely, plates with 0 mutants bring very little information (any sufficiently small mutation rate value would be a good fit for a dataset where most plates have no mutants). The experimental setup will thus always be tuned (for example through manipulation of population size or plating fraction) to avoid these two situations. With this rationale, when the dataset obtained through forward simulations of the fluctuation test with the parameters drawn as explained above had a median equal to 0 (more than half of the samples have no observed mutant) or higher than 1000, we reject it and start over the drawn of *µ* and *N*.

For simulations of fluctuation test with unknown fitness effect of the mutation, *f* was drawn uniformly over the range [0.4, 1.2]. This choice of range was motivated by both biological and practical reasons. Most mutations are deleterious, and the mutations classically used in a fluctuation test which provide resistance to antibiotics or other chemicals often incur a fitness cost in absence of the antibiotic. Moreover, a high fitness of the mutants increases the expansion of the mutant population and slows down the simulator, complicating the production of large test campaigns.

Death rate parameter *d* was uniformly drawn over the range [0, 0.7]. Values lower than 1 are necessary to correspond to the conditions of a fluctuation test, where population size increases to reach a large number of cells from a small inoculum. For the two-phase growth dynamics, both death rate values were independently drawn from this same distribution.

When it was not set to 1 (standard Luria & Delbrü ck model), plating efficiency (or plating fraction) *p* was uniformly drawn over the range [0.05, 0.25]. This aimed to represent the usual range of an intentional dilution when plating on selective medium. Intentional dilutions can in principle result in lower plating fractions (such as 0.001 in extreme cases); however, this implies a large number of mutants in the final culture before plating, which result in a slower simulation, hence for our large scale simulation study we did not consider such strong dilutions.

The initial number of cells *N*_0_ was always set to 1 except for simulations with cell mortality (*d >*0): in this case *N*_0_ was set to 100 to prevent a risk of population extinction. In experimental datasets, this number is higher than 1 for practical reasons (aiming for 1 cell per test tube in average will likely result in some of the test tubes being empty), but the exact value has no effect as long as it is low enough.

In most of the present work, we used our own simulator to produce the reference “experimental” datasets on which we benchmarked all methods. To ensure that this does not induce any bias in favor of our method, we also benchmarked all methods on datasets generated with the simulation tool proposed by rSalvador (Figure S24). We found that this does not affect the outcome.

### 5.4 Comparison with other methods

We compared the accuracy of our method with state-of-the-art tools based on analytical formulations of the probability distribution for the number of mutants.

#### rSalvador

We included rSalvador (Zheng, 2017), using the ‘`newton.joint.mk`’ function which implements the Mandelbrot-Koch model. This model allows to take into account differential fitness of the mutant (*f* ≠ 1) but also falls back to the ‘classical’ distribution (Lea and Coulson (1949) and Ma et al. (1992)) when *f* is set to 1. We thus did not explicitly include MSS-MLE (used by Falcor) in our benchmark. When *p* had to be provided to the estimator (Figures 5(a), 4(c)), we used the ‘`newton.LD.plating`’ function instead.

#### Flan

We also included Flan (Mazoyer et al., 2017), using the ‘`GF`’ method, which relies on the probability generating functions derived by Ycart (2014), based on a Bellman-Harris branching process. Bz-rates (Gillet-Markowska et al., 2015) implements the same methods (and makes them accessible through a web interface) and was thus not included in the comparison.

#### mlemur

Finally, we included mlemur, a very recent tool taking into account both differential fitness of the mutants and cell mortality (of the mutant as well as the wild type). We used the function ‘`mle.rate`’ of the mlemur R package.

Each method was run on each of the simulated datasets, and the parameter estimate returned by each tool was stored.

For ABC-MCMC, we retained the parameter(s)’s value(s) of the simulation presenting the best (lowest) KS score. We retained rSalvador and Flan’s estimated *µ* as the returned estimation of number of mutation events (m) divided by the total number of cells *N*. Mlemur’s estimated *µ* was directly returned by the ‘`mle.rate`’ function. rSalvador’s *f* estimation value was taken as returned from rSalvador’s estimation function (denoted as *w* in the function). Flan’s *f* estimation value was retrieved as 1 divided by the returned *ρ* value from the Flan’s estimation function (Flan’s *ρ* value is defined as the relative fitness of wild types compared to mutants, while in this paper, *f* is defined as the relative fitness of mutants compared to wild types).

Estimated parameter values were compared to the true values used as input of the simulation of the reference datasets. We report relative error (|(true_value-estimated_value)/true value|) for each parameter. The distributions of these estimation errors were compared for each tool. We tested statistically significant difference in the estimation error distributions between tools using Kruskal-Wallis tests followed by Dunn post-hoc tests.

## Supplementary materials

**Figure S1:**
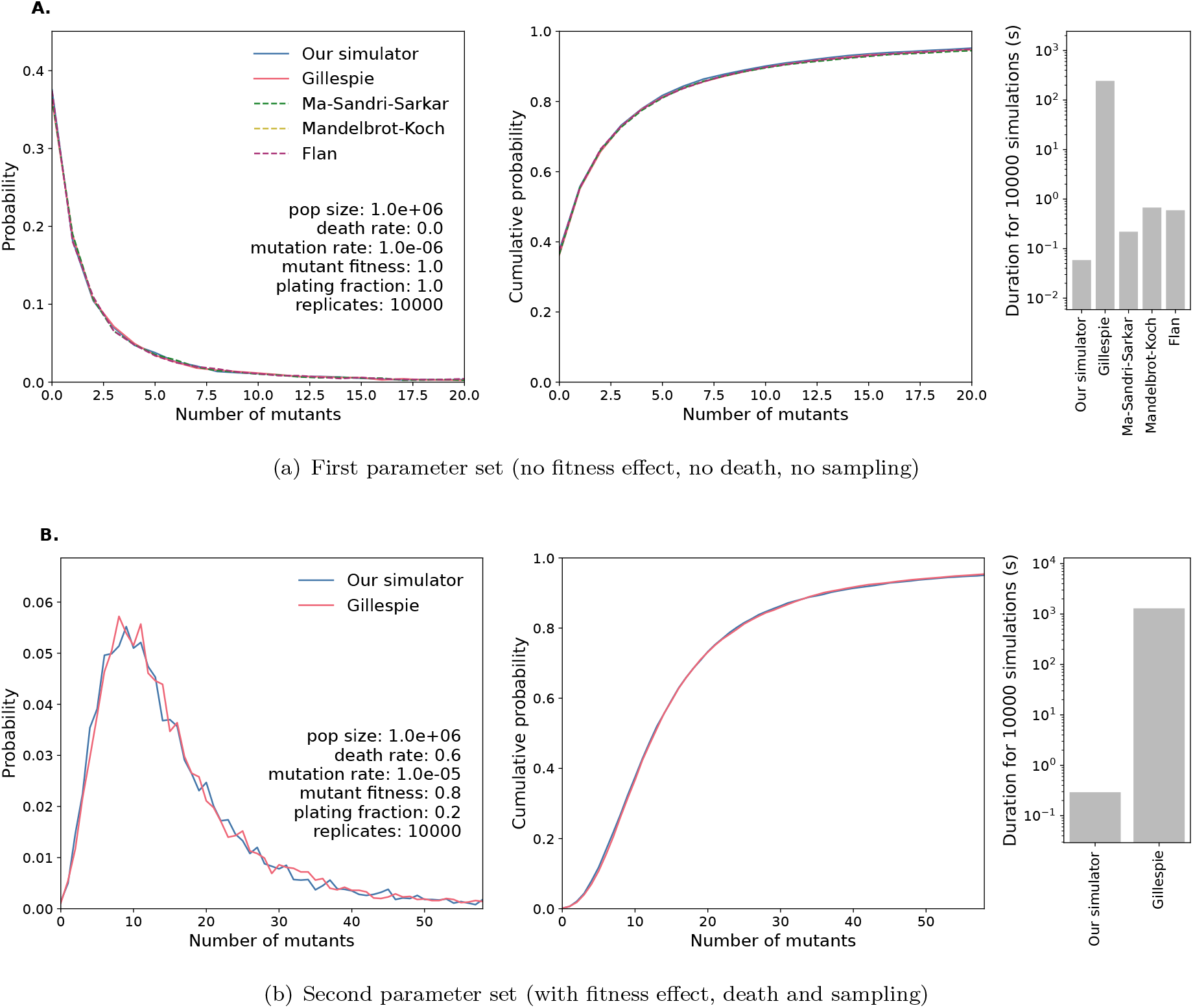
Test of the forward simulator, comparison with reference methods. For two parameter sets, the output of the simulator (10,000 simulated populations for each parameter set) is compared with the distributions obtained from the reference Gillespie simulator as well as thus obtained by other tools or theoretical models when suitable.

**Figure S2:**
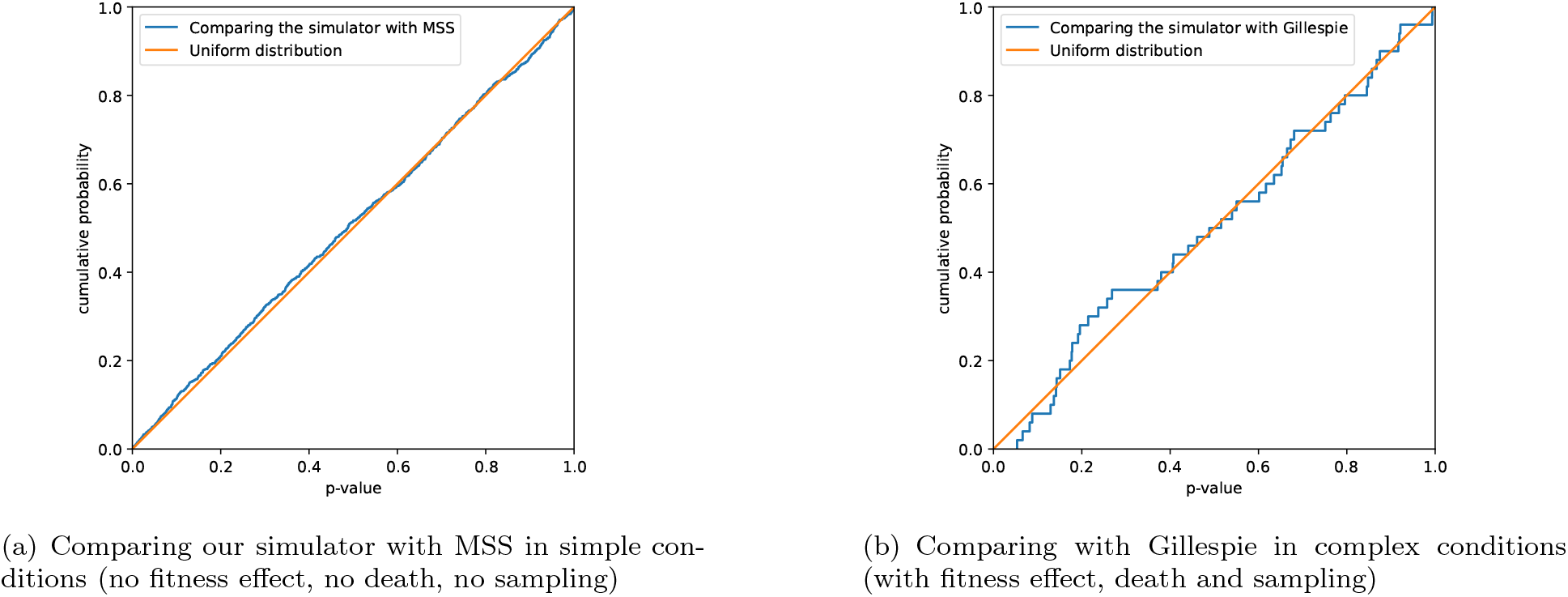
More thorough validation of the forward simulator: the previous comparison between our simulator and the state of the art is repeated on different parameter sets. **In panel (a)**, we compare the distribution obtained from our simulator with the distribution sampled from the Ma-Sandri-Sarkar (MSS) probability distribution in simple conditions (no fitness effect, no death, no sampling). The comparison is performed on 1,000 datasets, with randomly drawn parameters (population size and mutation rate). For each parameter set, we simulate 384 populations, and perform a two-sample Cramer-von Mises test to assess whether the obtained distribution differs from Ma-Sandri-Sarkar. As seen on the figure, the resulting p-values follow a uniform distribution over [0, 1], as expected if our simulator behaves exactly like MSS. **In panel (b)**, we perform the same comparisons between our simulator and simulations with the reference Gillespie algorithm; for 500 datasets with 96 replicate populations in complex conditions (fitness effect, death, and partial plating). Again, we find that the two methods produce undistinguishable distributions, as the p-values of the Cramer-von Mises tests are also uniformly distributed over [0, 1].

**Figure S3:**
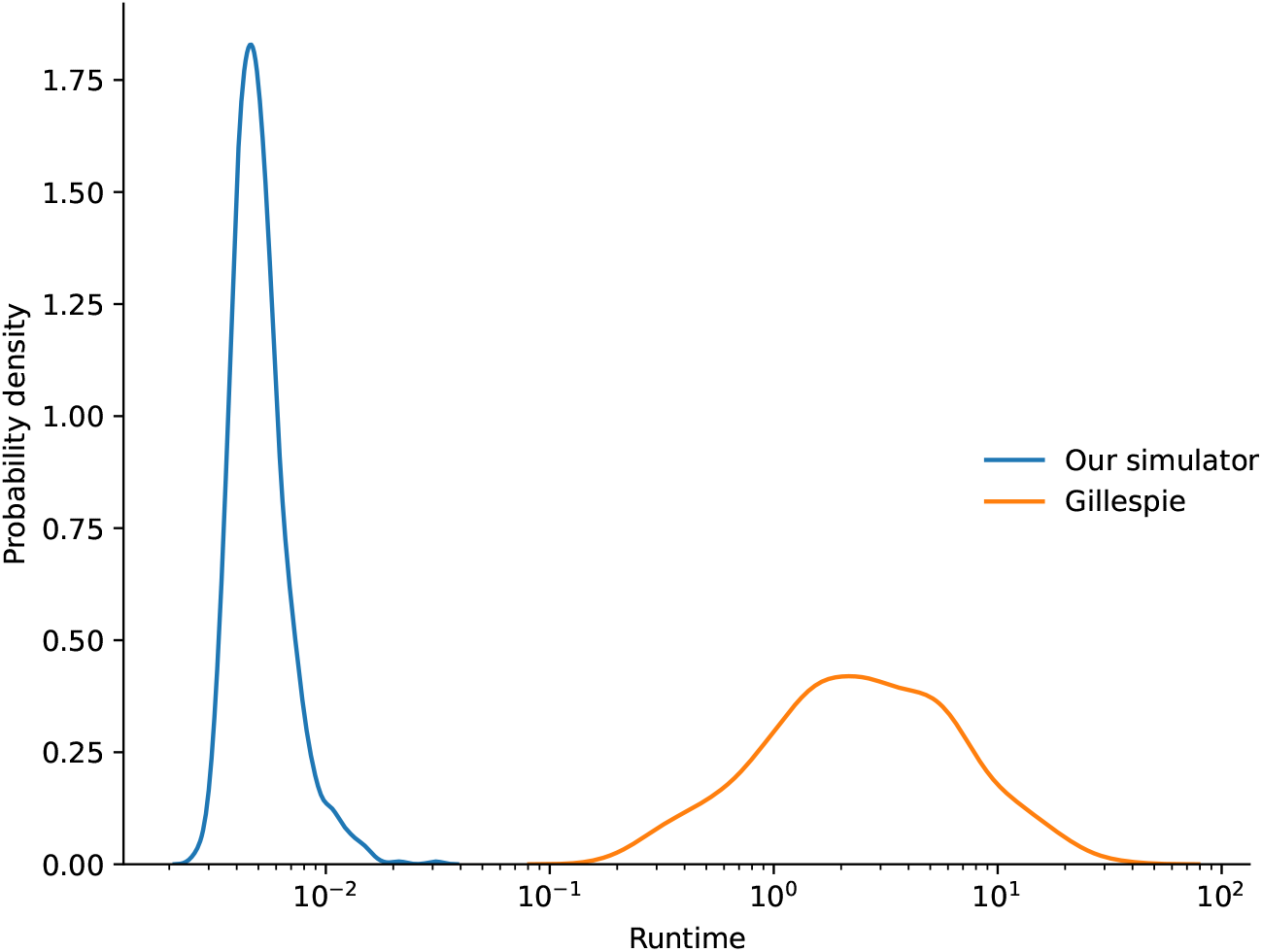
Comparison of the runtime of the forward simulator and Gillespie algorithm on a larger (1000) number of datasets with randomly drawn parameter values: the previous comparison between our simulator and the state of the art is repeated on 1000 randomly drawn parameter sets. For each parameter set, 384 populations were simulated. The average speedup provided by our simulator over Gillespie was a ratio 692 in run time.

**Figure S4:**
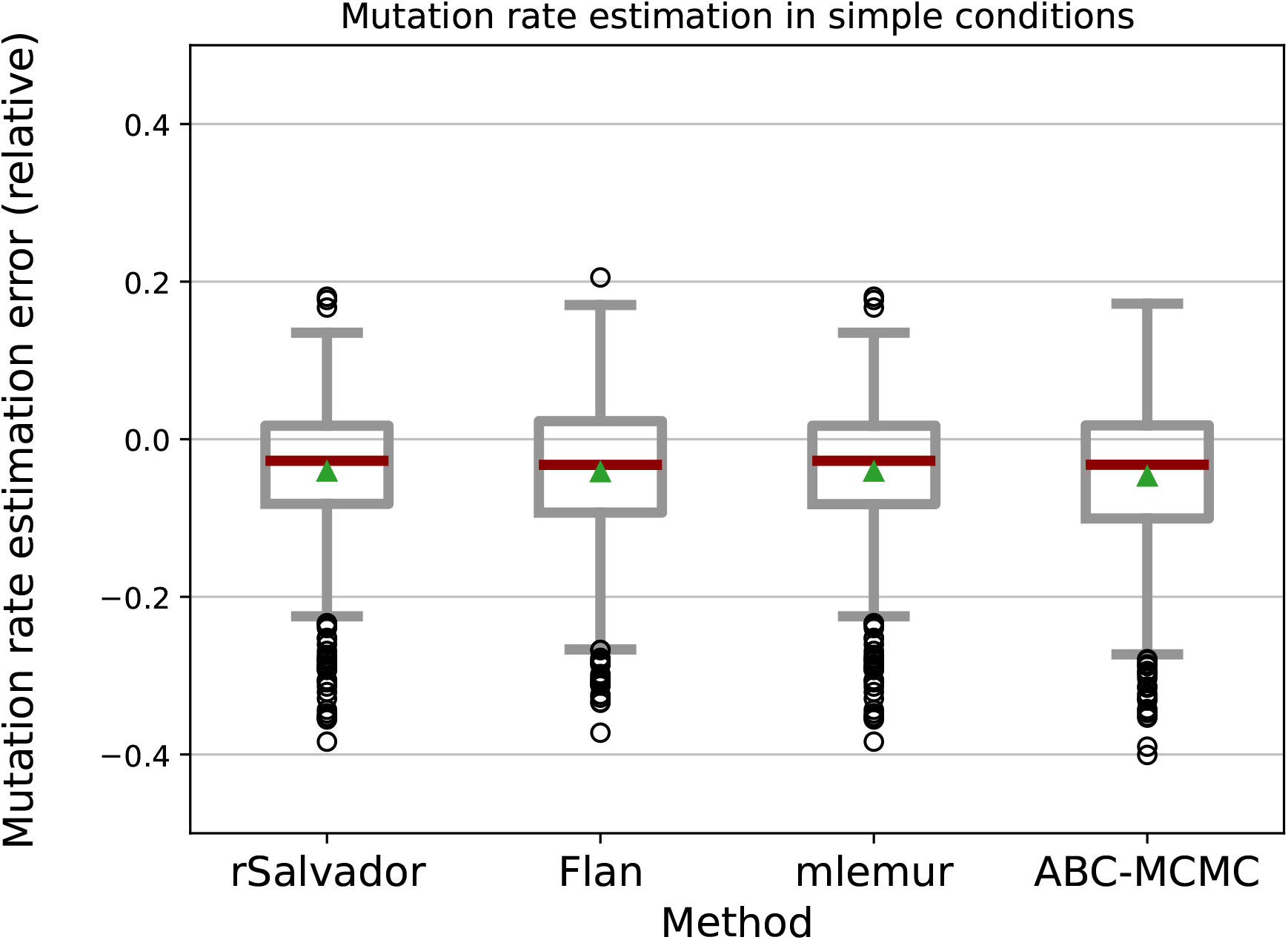
Mutation rate estimation errors in standard conditions, without absolute values. The estimation errors for all methods are shown for the same dataset than in Figure 3(a), but without taking the absolute value. The reported error is thus computed as ((true_value-estimated_value)/true_value). This reveals that all methods have a slightly more pronounced tendency to overestimate than to underestimate the mutation rate.

**Figure S5:**
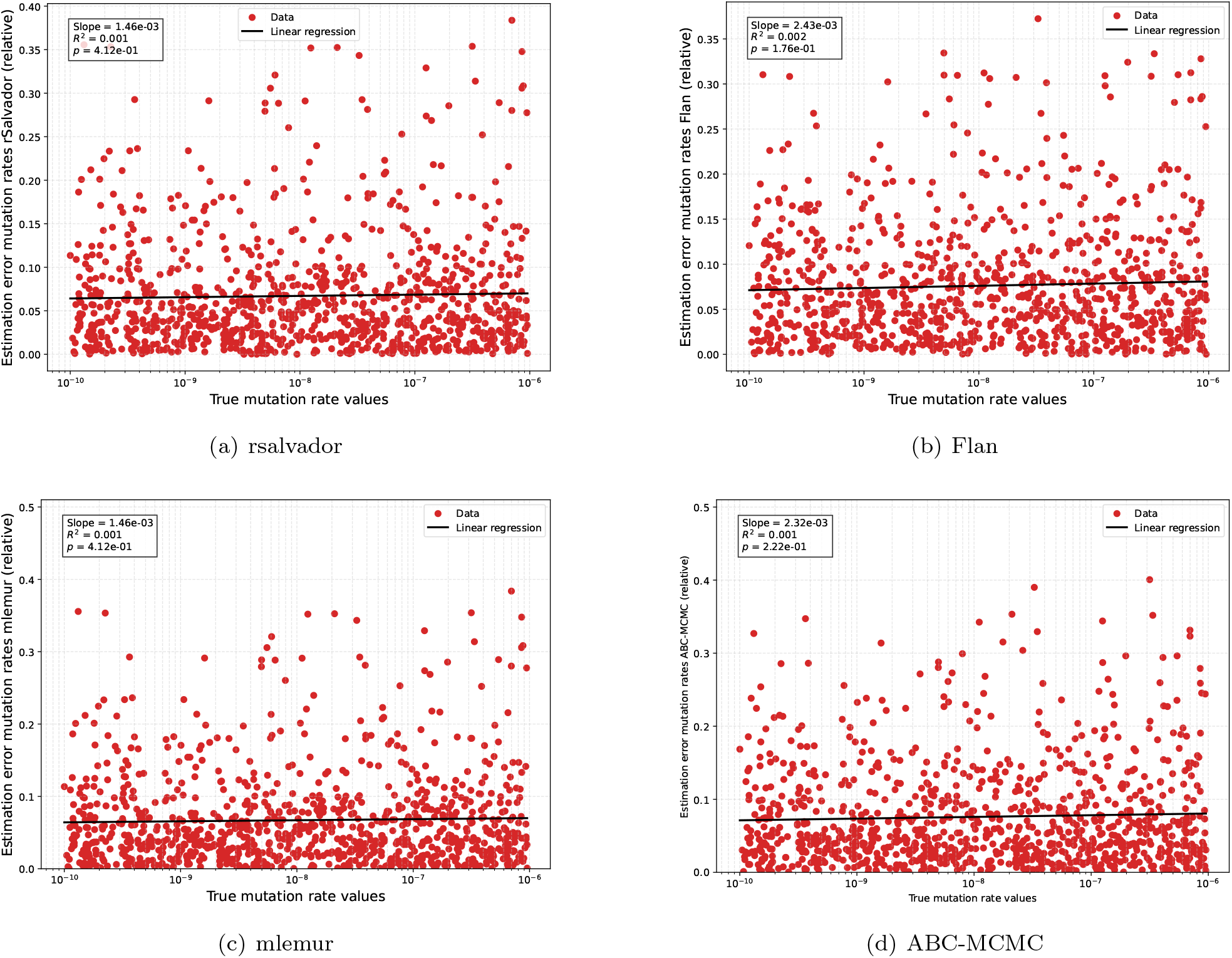
Mutation rate estimation errors as a function of the true mutation rate values. We show mutation rate (*µ*) estimation errors with rSalvador (a), Flan (b), mlemur (c), and ABC-MCMC (d), depending on the true *µ* values. *µ* is the only unknown parameter, other parameters were set to match the standard assumptions: no fitness difference between mutants and wild types (*f* =1), no death (*d*=0), full population plated without sampling and every mutant detected (*p*=1). Data were obtained using the same conditions as Figure 3(a), albeit with more replicates (1000 parameter sets, with 100 populations per parameter set). We found that the true mutation rate value does not affect the estimation error, which consistently remained around 0.05 over the full parameter range.

**Figure S6:**
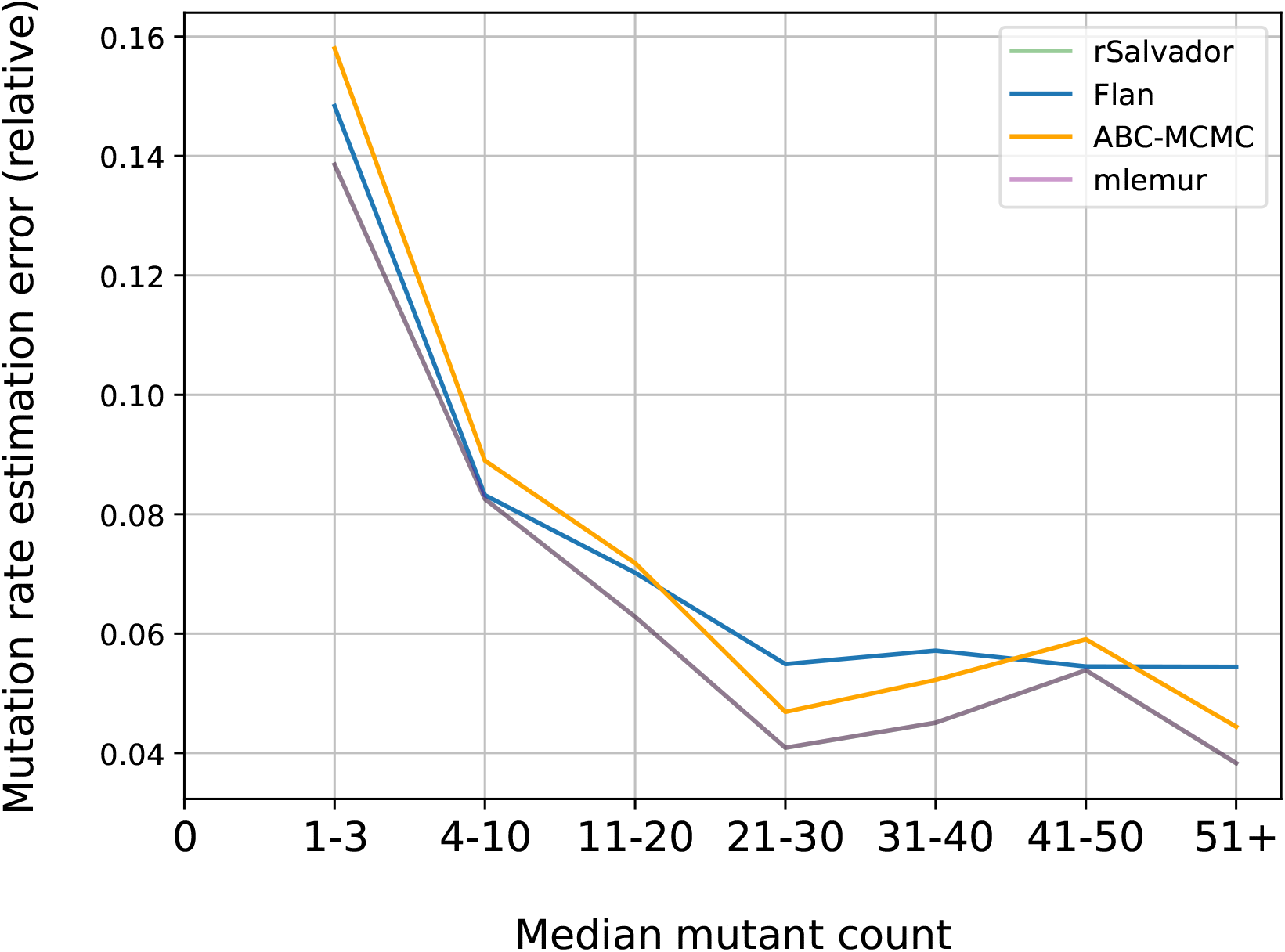
Effect of the median number of detected mutants on *µ* estimation error. Line plot of *µ* estimation error through rSalvador, Flan and ABC-MCMC depending on the median number of detected mutants in a fluctuation assay. We simulate 1000 fluctuation assay data sets with randomly drawn *µ* and N (see section 5.3) with *r* = 100 replicate populations. Other parameters were set to match the historical, standard assumptions (see Box 1): no fitness difference between mutants and wild types (*f* = 1), no death (*d* = 0), full population plated without sampling and every mutant detected (*p* = 1).

**Figure S7:**
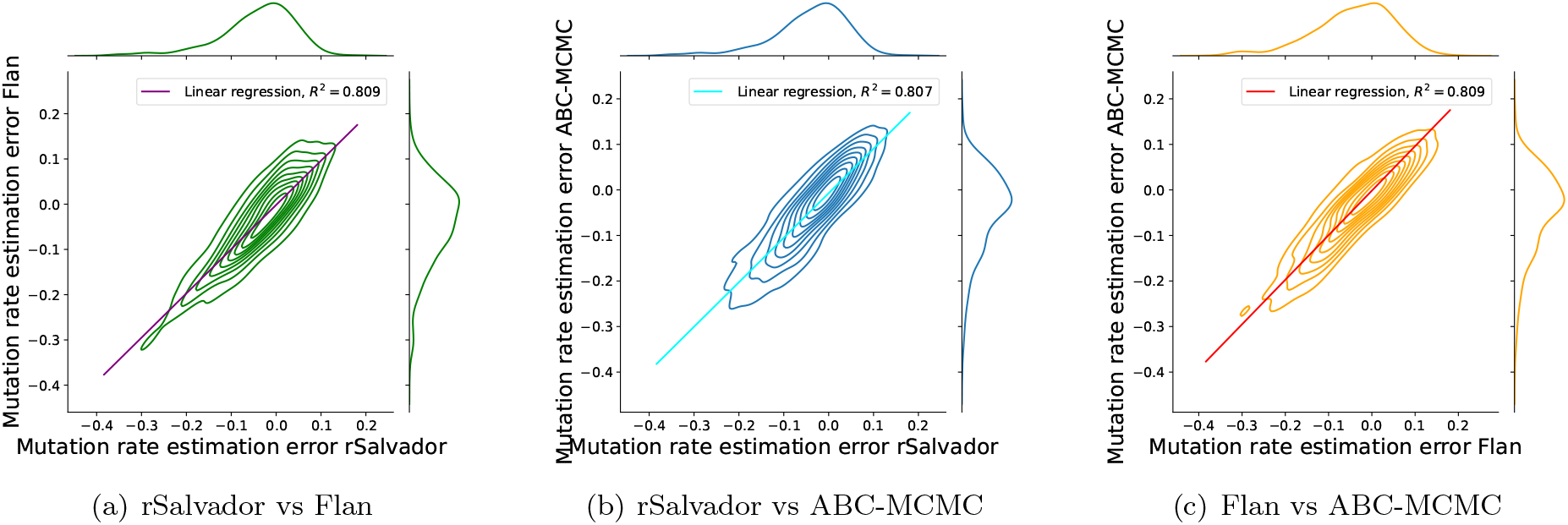
Joint plot of *µ* estimation errors between pairs of estimators. We compare *µ* estimation errors between different methods run on the same datasets. The dataset used for this figure was the same as in Figure S5. We used the error computed as computed as ((true_value-estimated_value)/true_value), without the absolute value. We found that datasets that are difficult to accurately estimate for a method are also difficult to estimate for the others methods, with highly correlated errors between methods (*R*^2^ values are 0.809, 0.807 and 0.809 for panels a, b, and c respectively). Contour lines represent the kernel density estimates of the joint probability distributions: each line represents an isoprobability, and 10 different lines are draw at 10 equally spaced cumulative probabilities (the inner contour line contains 10% of the data points in a smoothed model, the next one 20%, then 30%, etc).

**Figure S8:**
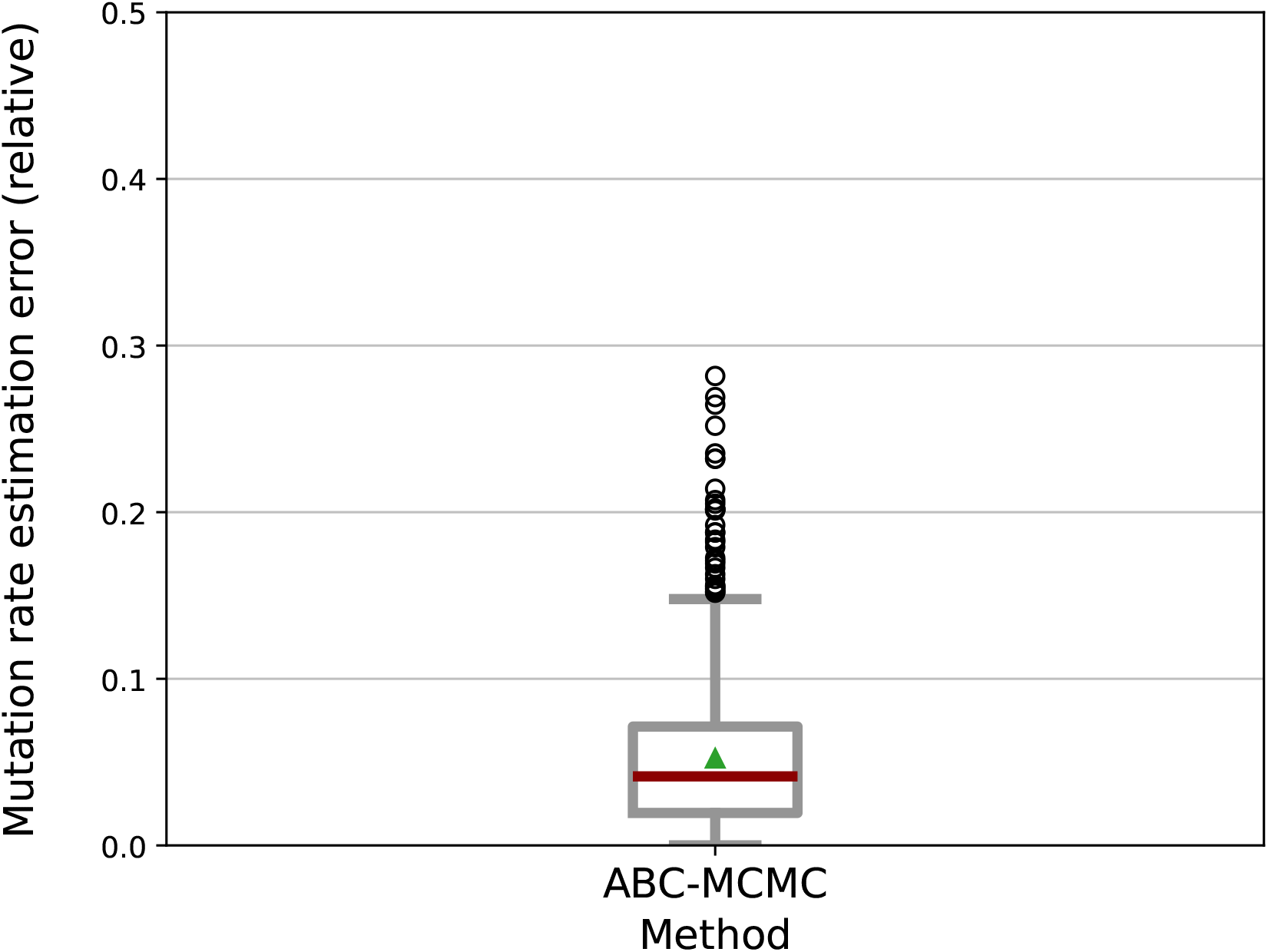
Mutation rate estimation with known, piecewise constant death, known fitness effect of the mutation and sampling at a known fraction. This represents the same scenario as in Figure 5b, with the addition of a known fitness effect of the mutation (*f* ≠ 1) and sampling at a known rate (*p* ≠ 0).

**Figure S9:**
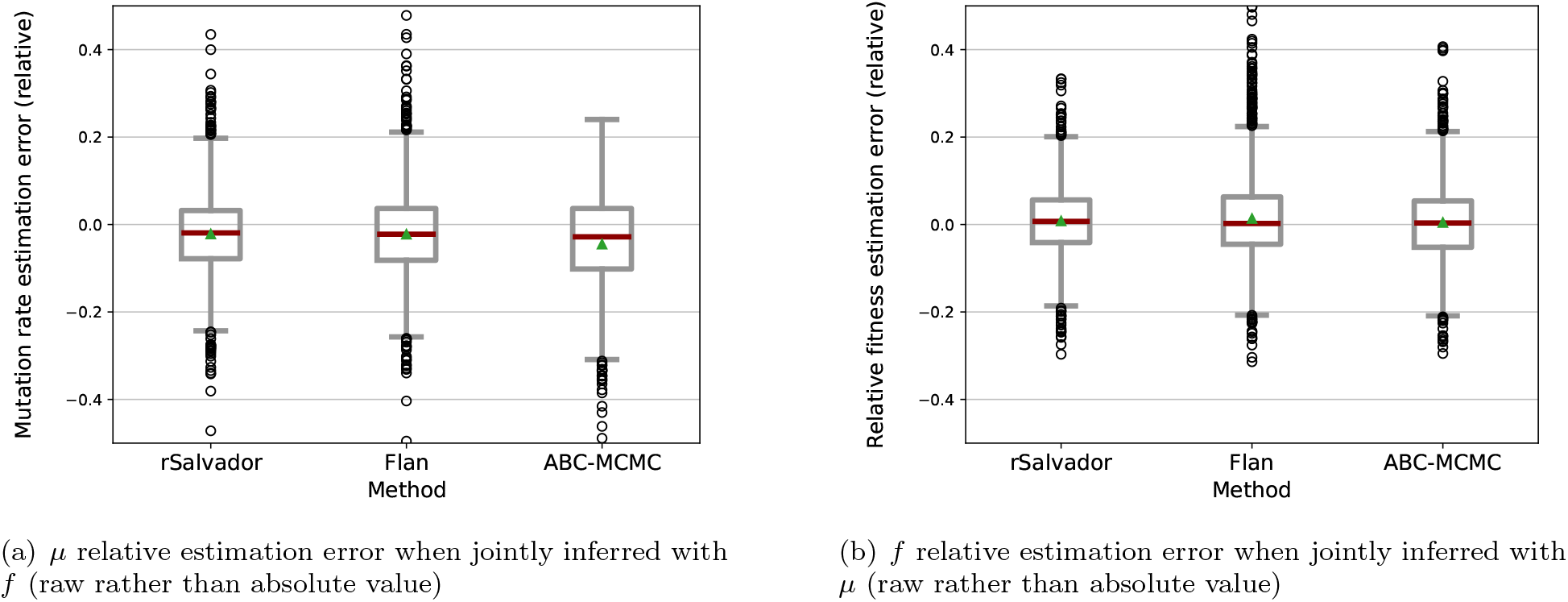
Joint *µ* and *f* estimation, with relative error reported without absolute value. The data set is the same than for Figure 6, but relative estimation errors are reported with raw rather than absolute values. The reported error is thus computed as ((true value-estimated value)/true value). As in Figure S4, we find that all methods have a slightly more pronounced tendency to overestimate than to underestimate the mutation rate, but show no bias when estimating the fitness effect.

**Figure S10:**
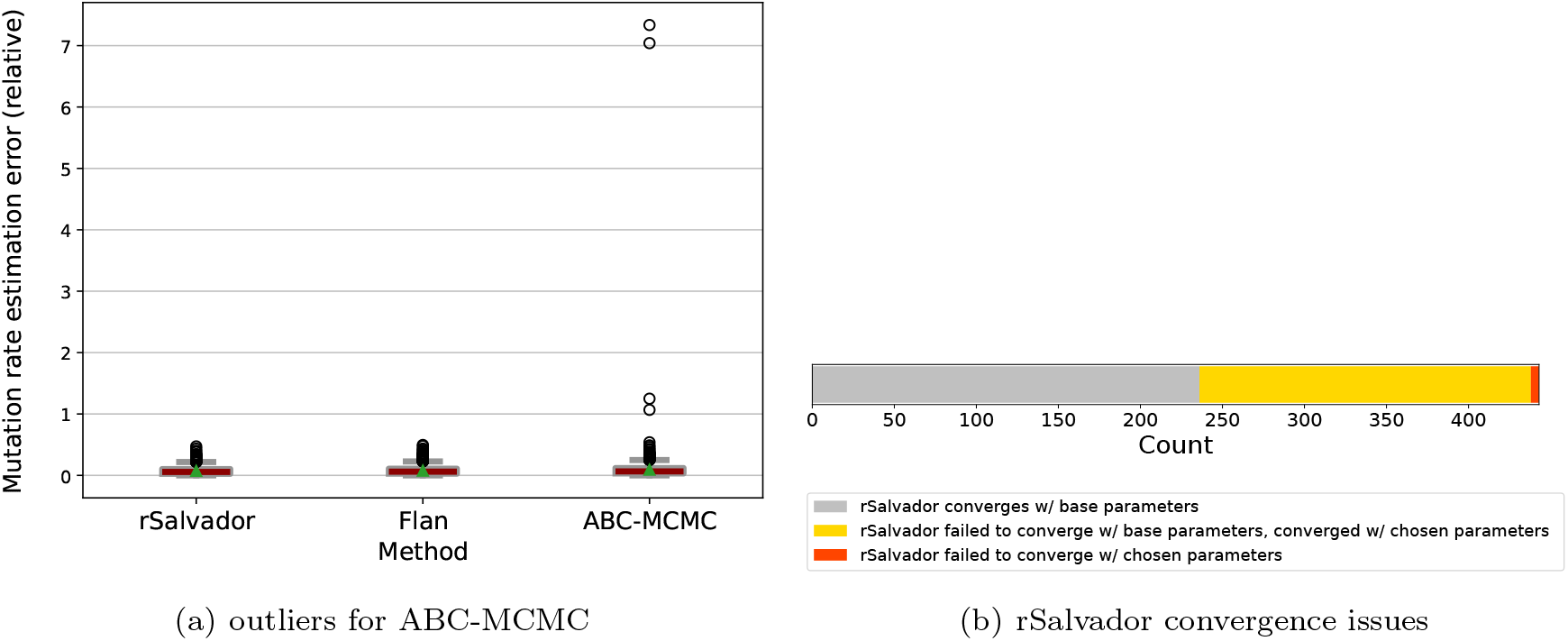
Joint *µ* and *f* estimation: outliers and convergence issues. The same dataset is used than in Figure 6, but on panel (a) the zoom level on the y axis is chosen to show the outliers, and on panel (b) convergence issues with rSalvador are shown. **(a)** As a stochastic parameter search method itself based on stochastic simulations, ABC-MCMC is prone to rare but strong outliers. These outliers were included in the quartiles shown in Figure 6. **(b)** rSalvador joint estimation of *µ* and *f* sometimes fails to converge when the function is called with default parameters. For 443 simulated datasets, 236 (53%) converged without error using the base parameters (starting point of the Newton-Raphson search). 202 (46%) failed to converge using the base parameters, but successfully converged after changing the starting point parameters. 5 (1%) kept failing even after several attempts of changing the starting point parameters. We used a predefined list of starting point values (chosen purely empirically by manually testing on a few examples), which were tested sequentially until the algorithm converged. The cases where rSalvador failed to converge with default parameters but converged with specific parameters were included in Figure 6, but those where it did not converge in any of our attempts were excluded from this dataset, not only for rSalvador but also for the other tools.

**Figure S11:**
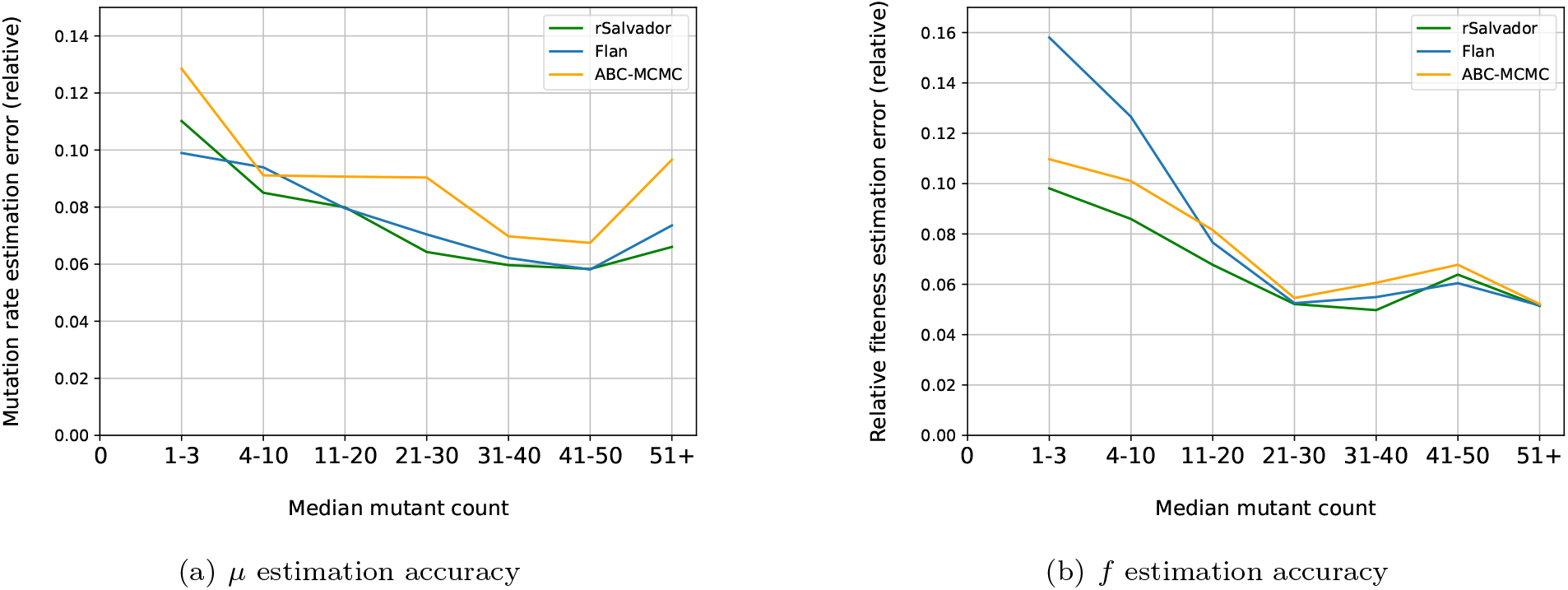
Effect of the median number of detected mutants on joint *µ* and *f* estimation errors. Panels (a) and (b) show estimation errors of *µ* and *f* through rSalvador, Flan and ABC-MCMC depending on the median number of detected mutants. We use the same 1943 simulated fluctuation assays than in Figure 6.

**Figure S12:**
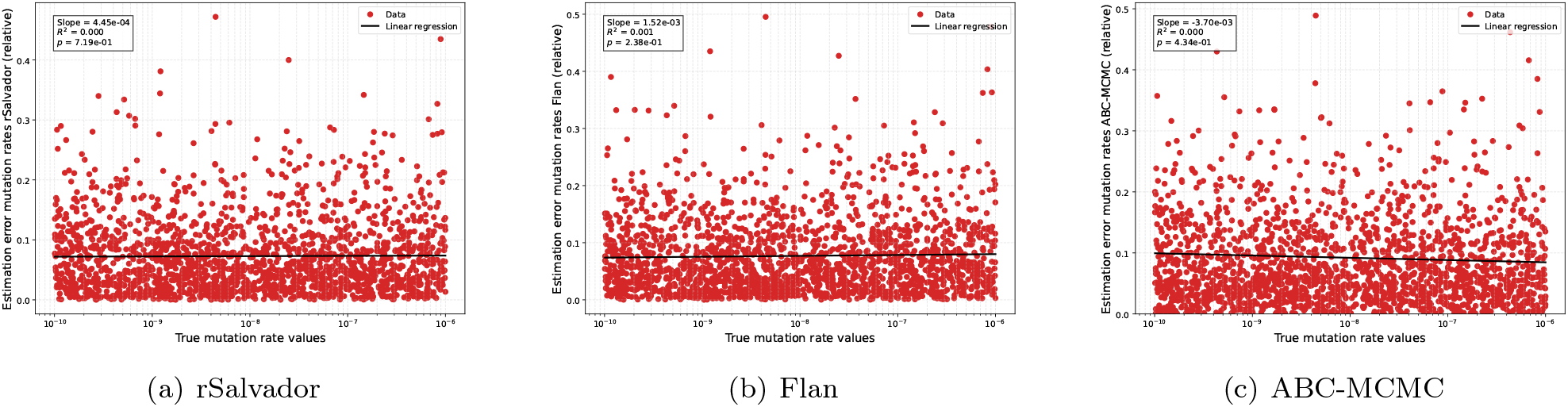
Mutation rate estimation errors as a function of the true mutation rate value when the mutation rate and the fitness effect are simultaneously inferred. The plots show the mutation rate (*µ*) estimation errors obtained with rSalvador (a), Flan (b) and ABC-MCMC (c) depending on the true *µ* values. We simulated fluctuation assays with mutation rate *µ, f* and N randomly drawn, and jointly inferred *µ* and *f*. Other parameters were set to match the historical, standard assumptions (death rate *d* = 0, sampling fraction *p* = 1). The dataset used for this figure was the same as in Figure 6. As when inferring *µ* alone (Figure S5), we found no link between the true value and the estimation error.

**Figure S13:**
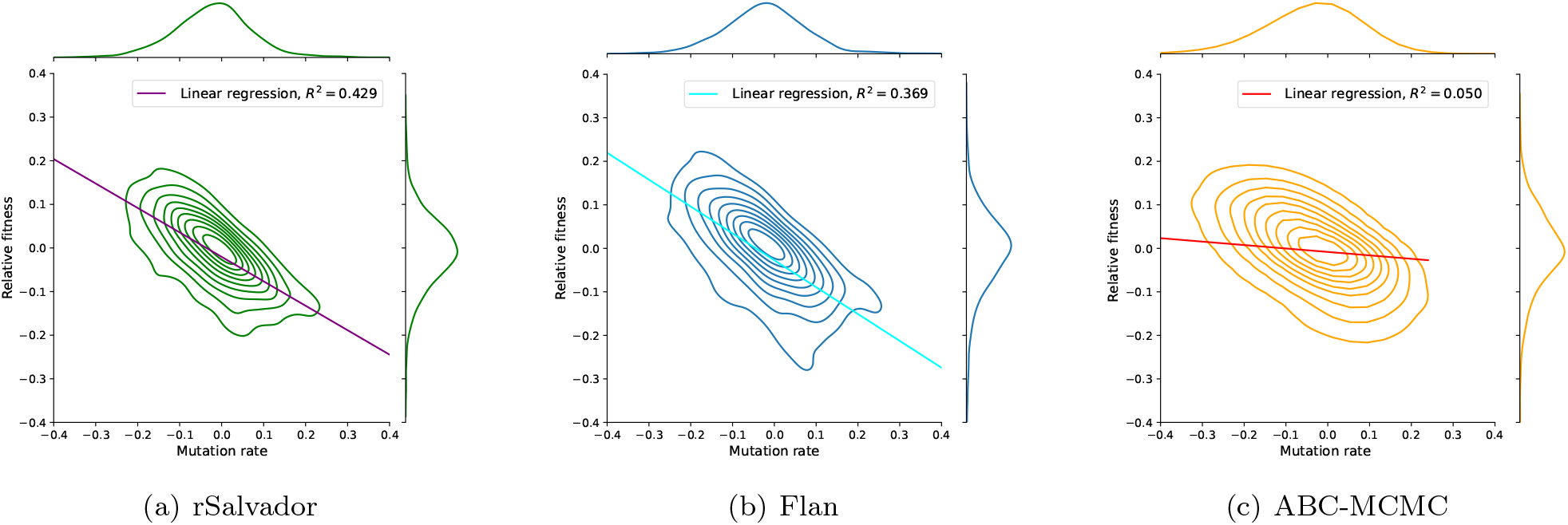
Joint distributions for *µ* and *f* estimation errors when simultaneously inferred. The axes represent the relative estimation error values, without the absolute values, computed as ((true value-estimated value)/true value). Contour lines represent the kernel density estimates of the joint probability distributions, as in Figure S7. The dataset used for this figure was the same as in Figure 6. Estimation errors between the two parameters are more strongly correlated for rSalvador and Flan than for ABC-MCMC (*R*^2^ values are 0.429, 0.369, and 0.050, respectively), but all correlations are significant.

**Figure S14:**
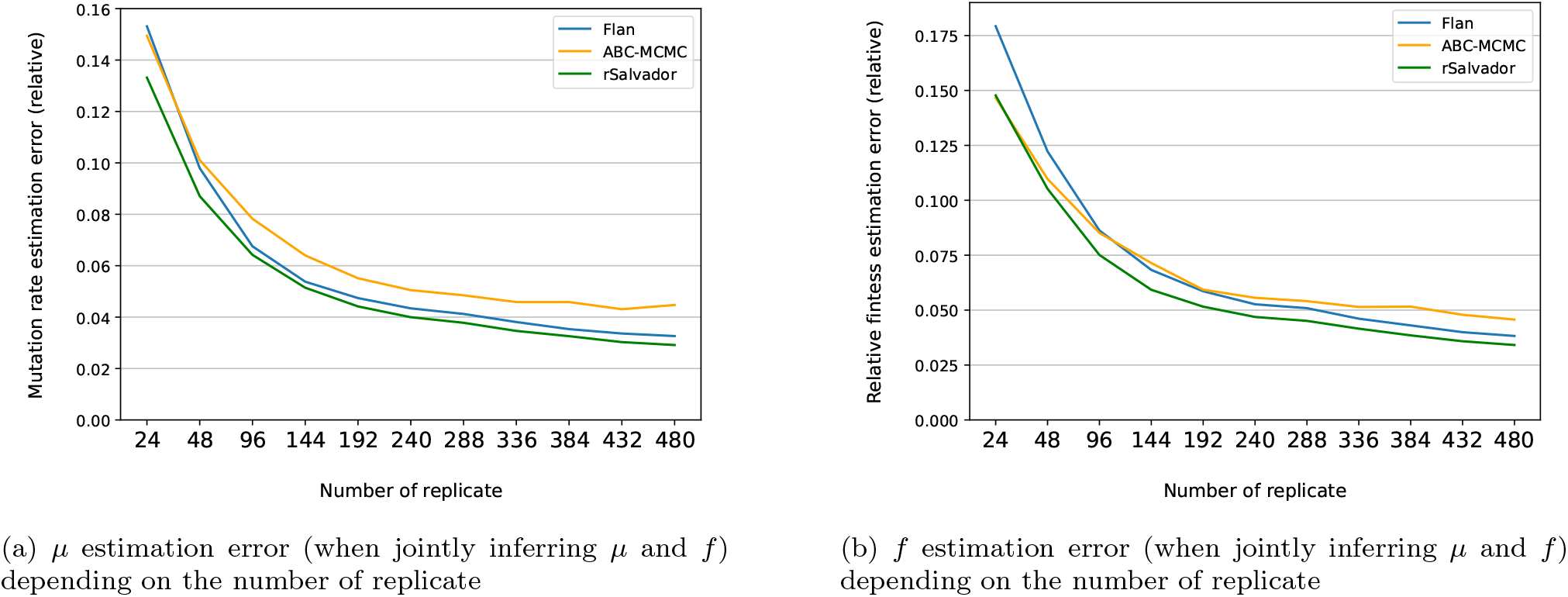
Effect of the number of replicate populations on the *µ* and *f* joint estimation errors. Line plot of *µ* (panel **a**) and *f* (panel **b**) estimation errors obtained with rSalvador, Flan and ABC-MCMC,depending on the number of replicate populations in a fluctuation assay. We simulate 438 fluctuation assays with *µ, f* and *N* randomly drawn, with *r* = 480 replicate populations, and jointly estimate *µ* and *f* when subsets of a given size of the mutant counts are given to the estimator. Other parameters were set to match the standard assumptions (death rate *d* = 0, plating fraction *p* = 1).

**Figure S15:**
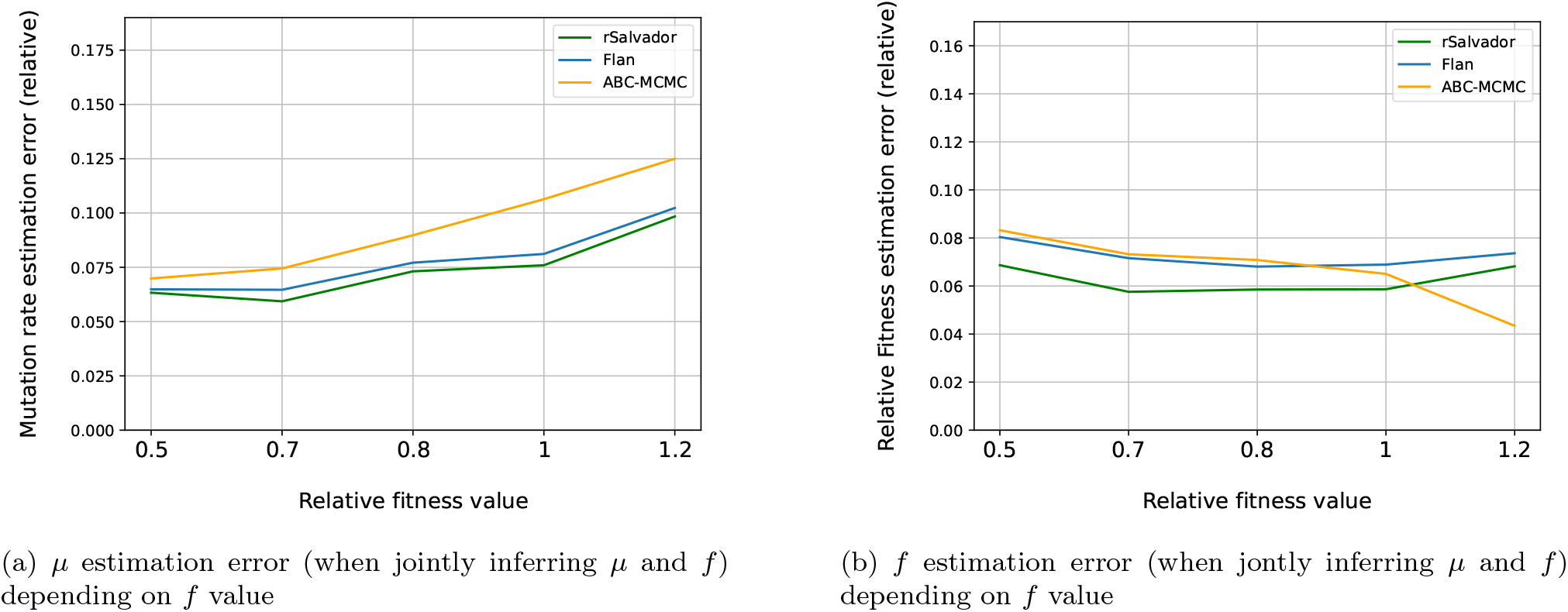
Effect of *f* value on the joint *µ* and *f* estimation accuracy. Line plot of *µ* (panel **a**) and *f* (panel **b**) estimation errors through rSalvador, Flan and ABC-MCMC depending on datasets *f* value. We used the same 1943 simulated fluctuation assays than in Figure 6. The results from panel (b) are shown in more details in Figure S16

**Figure S16:**
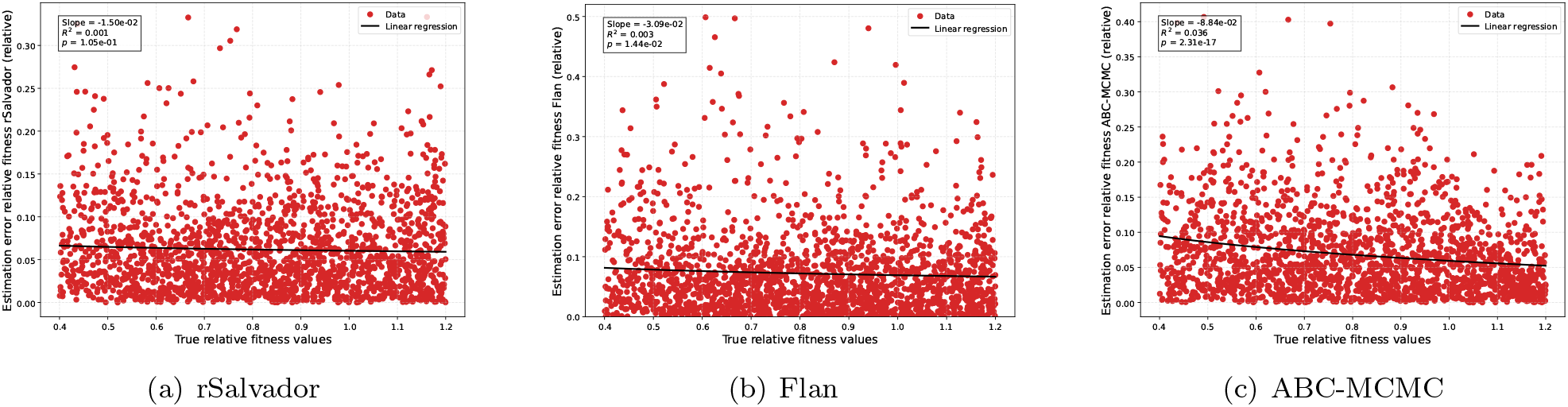
Relative fitness estimation errors as a function of the true relative fitness values. Relative fitness (*f*) estimation errors obtained with rSalvador (a), Flan (b) and ABC-MCMC (c), depending on the true *f* values. We simulated fluctuation assays with mutation rate *µ, f* and N randomly drawn, and jointly inferred *µ* and *f*. Other parameters were set to match the historical, standard assumptions (death rate *d* = 0, sampling fraction *p* = 1). The dataset used for this figure was the same as in Figure 6. Compared to panel (b) of Figure S15, we here display all individual data points. We found that the true fitness value is negatively correlated with the fitness estimation error for Flan (very weak correlation) and ABC-MCMC (stronger correlation), but not for rSalvador.

**Figure S17:**
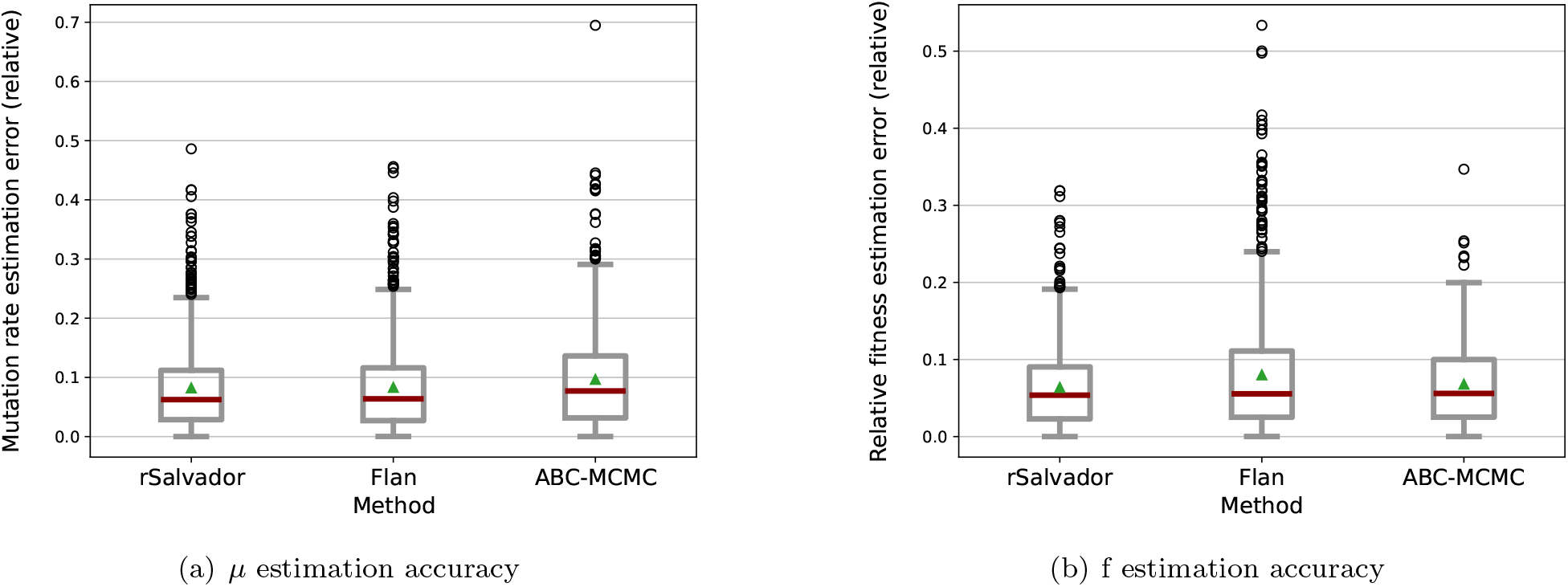
Joint *µ* and *f* estimation in standard conditions, when *f* = 1 but the estimator does not know this and attempts to infer it. Boxplots of the overall *µ* estimation accuracy (a) and *f* accuracy (b) through rSalvador, Flan and ABC-MCMC with 2 unknown parameters (*µ* and *f*). We simulated 880 fluctuation assay data sets with *µ* and N randomly drawn with *r* = 100 replicate populations. *f* was always set to 1, but the methods attempted to infer it jointly with *µ*. Other parameters were set to match the historical, standard assumptions: no death (*d* = 0), full population plated without sampling and every mutant detected (*p* = 1).

**Figure S18:**
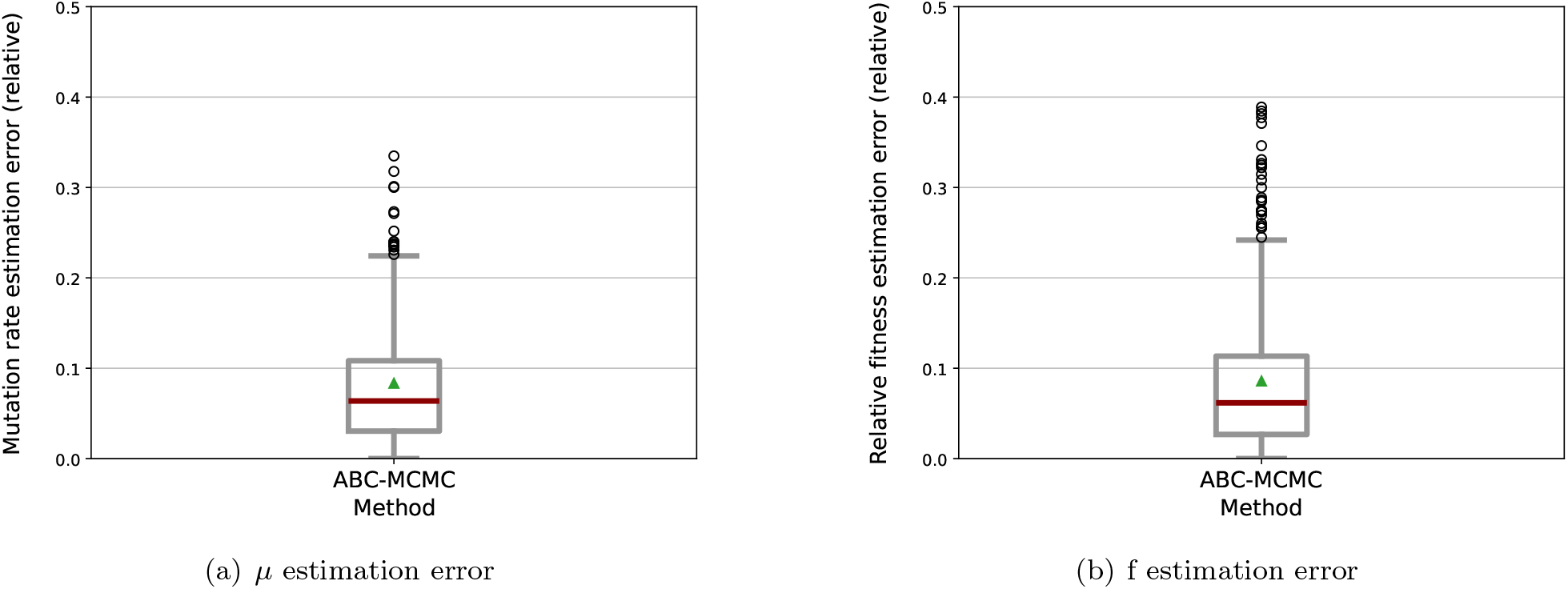
Joint *µ* and *f* inference in non-standard conditions: known cell mortality (*d* ≠ 0). We simulated 550 fluctuation assay data sets with *µ, f, d* and *N* randomly drawn with *r* = 100 replicate populations, and jointly inferred *µ* and *f* with ABC-MCMC, giving the known *d* value to the estimator. Other parameters were set to match the historical, standard assumptions (plating fraction *p* = 1).

**Figure S19:**
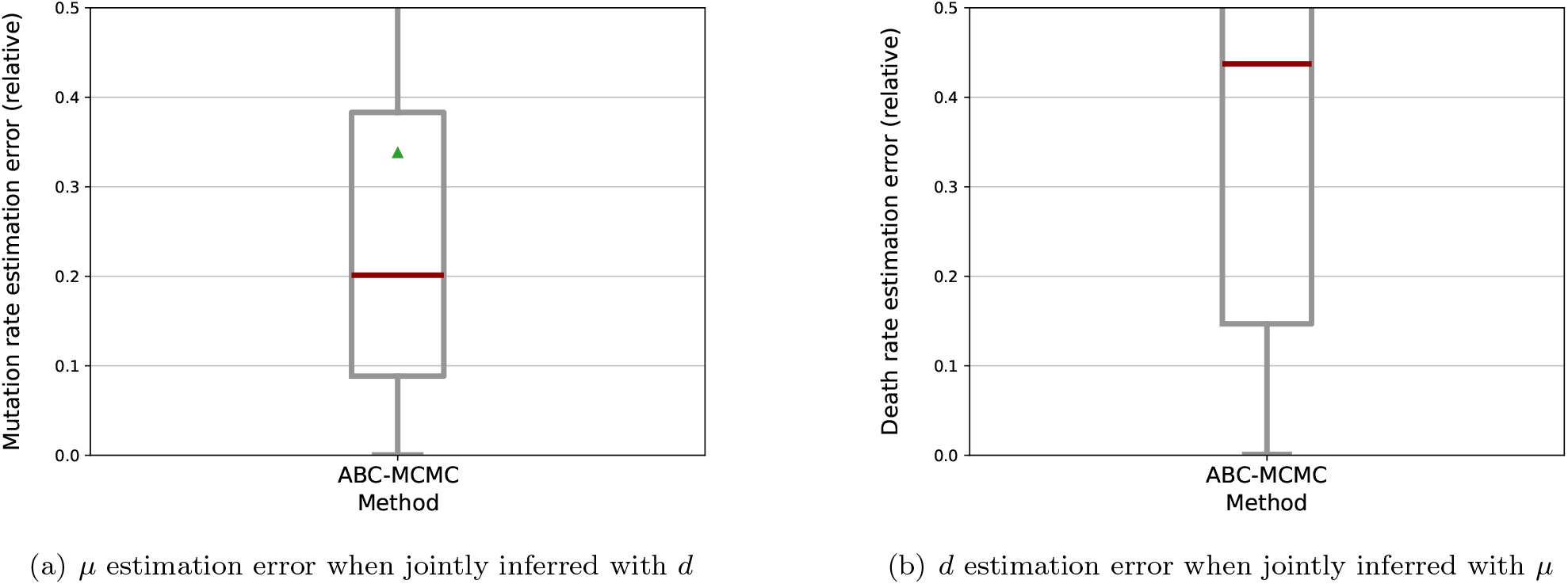
Joint inference of *µ* and *d* with ABC-MCMC. We simulated 370 fluctuation assay data sets with *µ, d* and *N* randomly drawn, with *r* = 100 replicate populations, and jointly inferred *µ* a *d*. Other parameters were set to match the historical, standard assumptions (relative fitness *f* =1, sampling fraction p=1).

**Figure S20:**
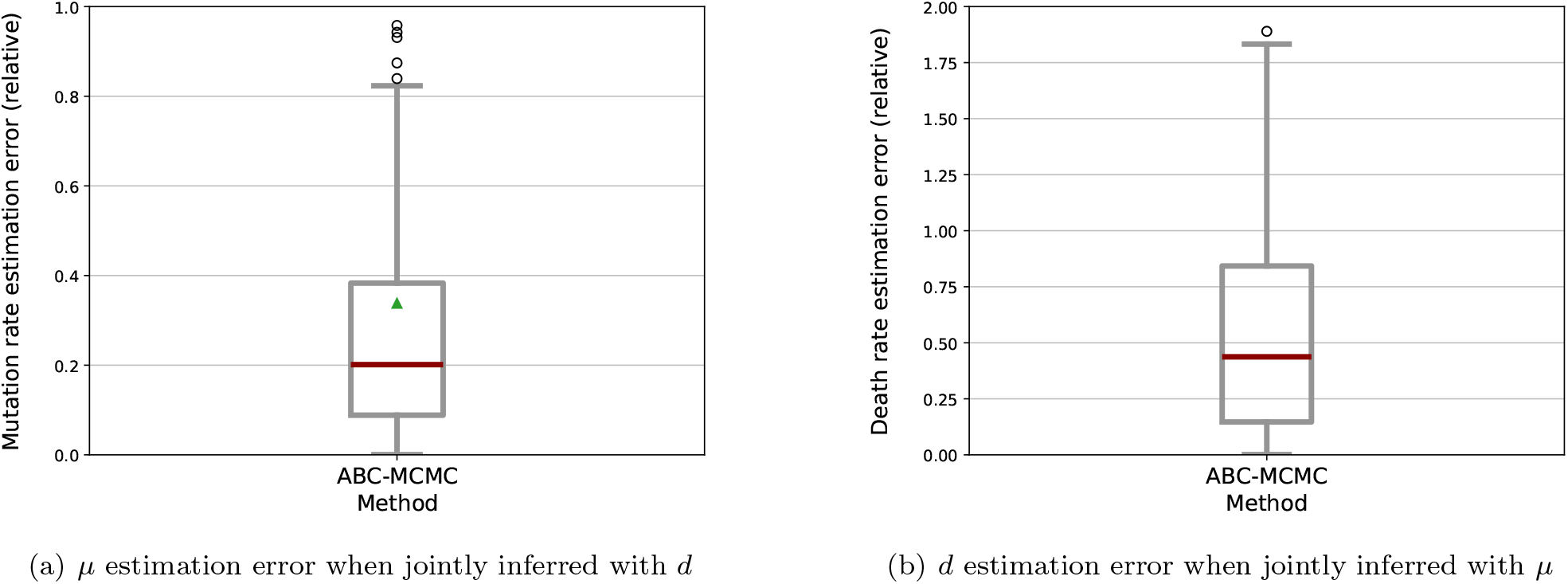
Joint inference of *µ* and *d* with ABC-MCMC with extended y-axes and full boxplot shown. We simulated 370 fluctuation assay data sets with *µ, d* and *N* randomly drawn, with *r* = 100 replicate populations, and jointly inferred *µ* a *d*. Other parameters were set to match the historical, standard assumptions (relative fitness *f* =1, sampling fraction p=1).

**Figure S21:**
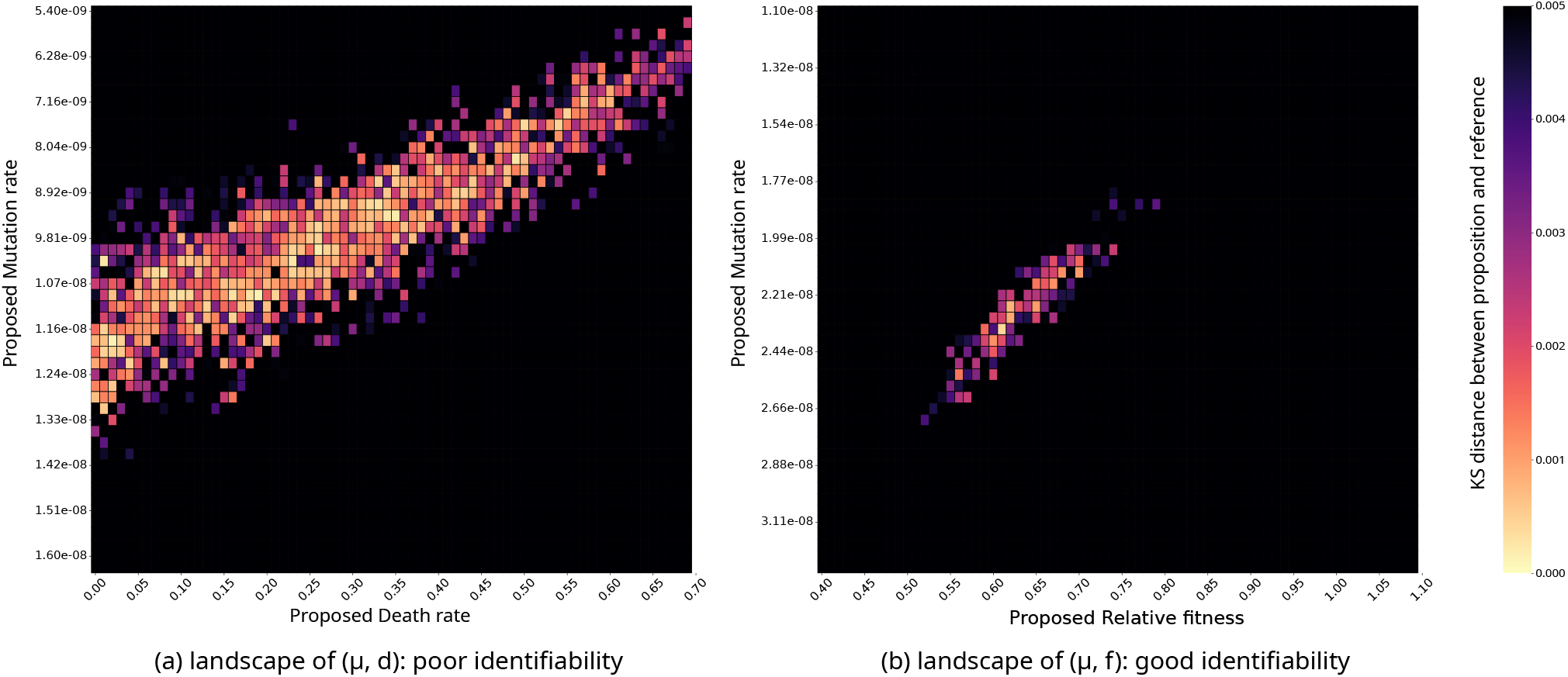
Parameter fit landscape for joint estimations of (a) mutation rate and death rate or (b) mutation rate and relative fitness. Distributions obtained for all exhaustive combinations of parameter pairs are compared to a reference distribution (obtained with random values for N, *µ* and *f* or *d*). *r* = 500 populations are simulated for each parameter combination as well as for the reference simulation. The heatmap shows for each parameter pair the distance (KS score) between the distribution of the number of mutants obtained when simulating with the parameter pair and the reference distribution. Brighter colors indicate lower distances. True values used for the reference datasets were (a) *µ* = 1.08 *×* 10^*−*8^ and *d* = 0.16, (b) *µ* = 2.2 *×* 10^*−*8^ and *f* = 0.66.

**Figure S22:**
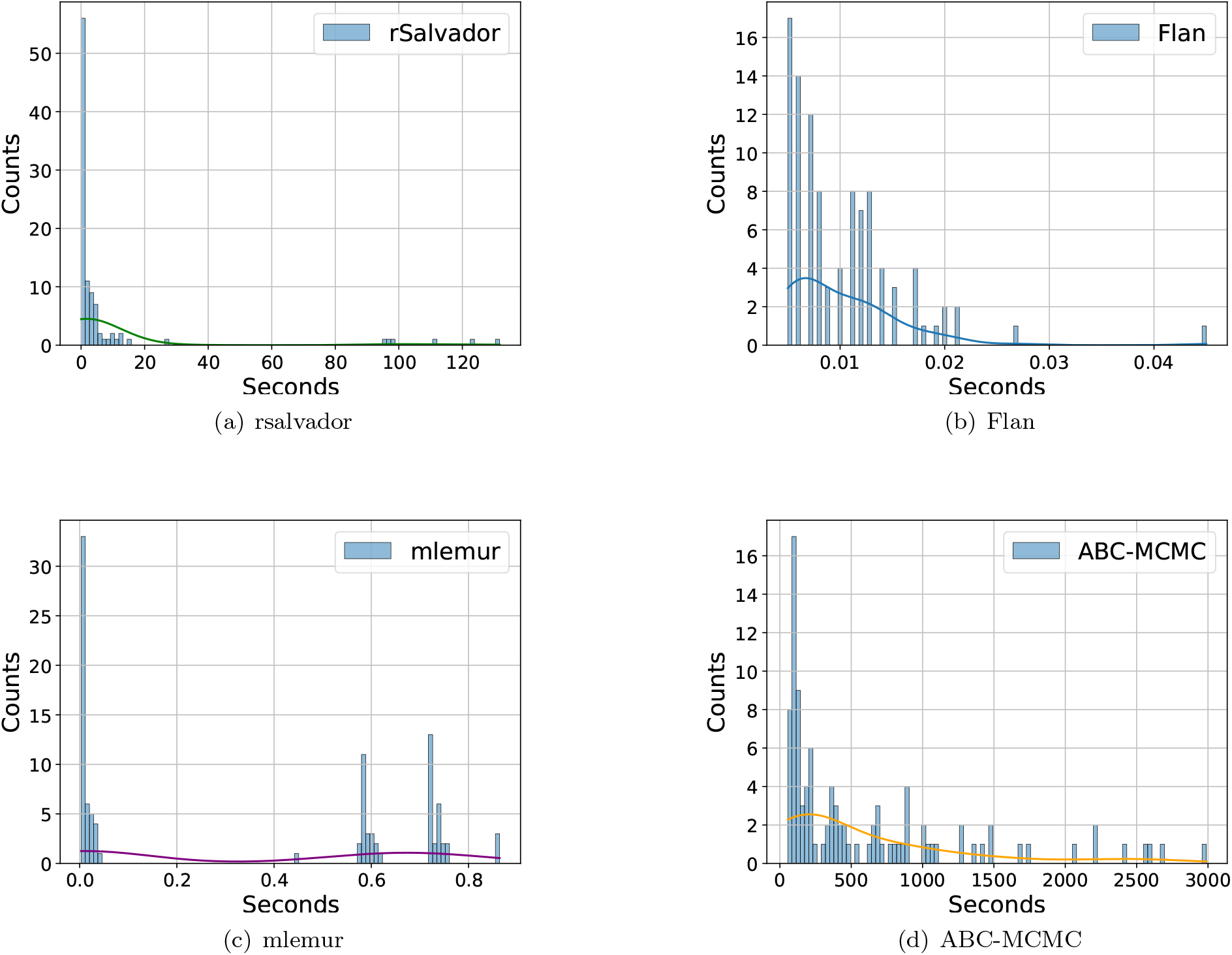
Running time of *µ* inference for rSalvador, Flan, mlemur and ABC-MCMC in classical conditions. For each tool, we represented the probability distribution (over 100 datasets) of the computation time taken to perform *µ* inference on one dataset. The curves represent smoothing by a kernel density estimation. *µ* was the only unknown parameter, other parameters were set to match the standard assumptions: no fitness difference between mutants and wild types (*f* =1), no death (*d*=0), full population plated without sampling and every mutant detected (*p*=1). For each simulated dataset, *µ* was randomly drawn from a log-uniform distribution over the range [10^*−*10^, 10^*−*6^], *N* was subsequently randomly drawn from a log-uniform distribution over the range [10^*−*2^, 10^2^]*/µ*. It should be noted that the choice of *N* and *µ* values drastically affects the efficiency of both simulators. Gillespie algorithm becomes slower when *N* becomes large. The speed of our simulator rather depends on the number of mutants, so on the product *N × µ*.

**Figure S23:**
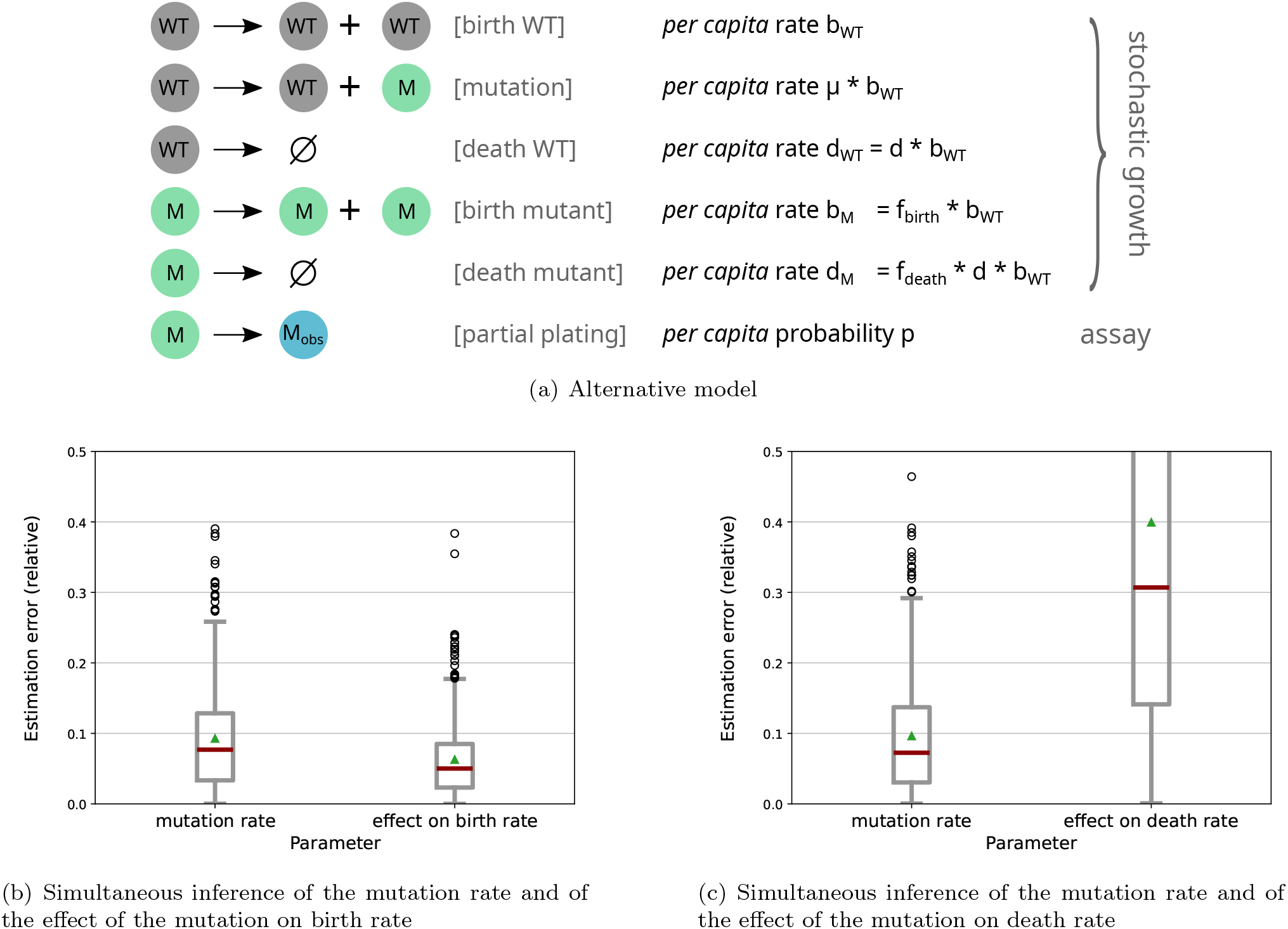
Alternative model with independent effects of the mutation on birth rate and on death rate. In the main model presented in the article, death rate is proportional to birth rate (the death parameter is the rate of death compared to the rate of birth), and thus the mutation has the same multiplicative effect on birth rate and death rate of the mutant. In this alternative model, the mutation can have independent effects on the mutant birth rate and the mutant death rate. This effect is thus characterized by two parameters (*f*_*birth*_ and *f*_*death*_) instead of one (*f*). This alternative model is schematized in panel (a). We successfully tested the ability of ABC-MCMC to jointly infer the mutation rate and the effect of the mutation on birth rate, as presented in panel (b). This was done in the presence of death (with a known death rate and a known effect of the mutation on this death rate), in absence of sampling before plating. 571 fluctuation test assays were simulated, with parameters randomly drawn in the same range than for other figures. Each simulation assay had *r* = 96 replicate populations. We also show (panel (c)) that ABC-MCMC fails to jointly infer the mutation rate and the effect of the mutation on death rate, consistent with the inability to jointly infer the mutation rate and death rate in the main model presented in the article.

**Figure S24:**
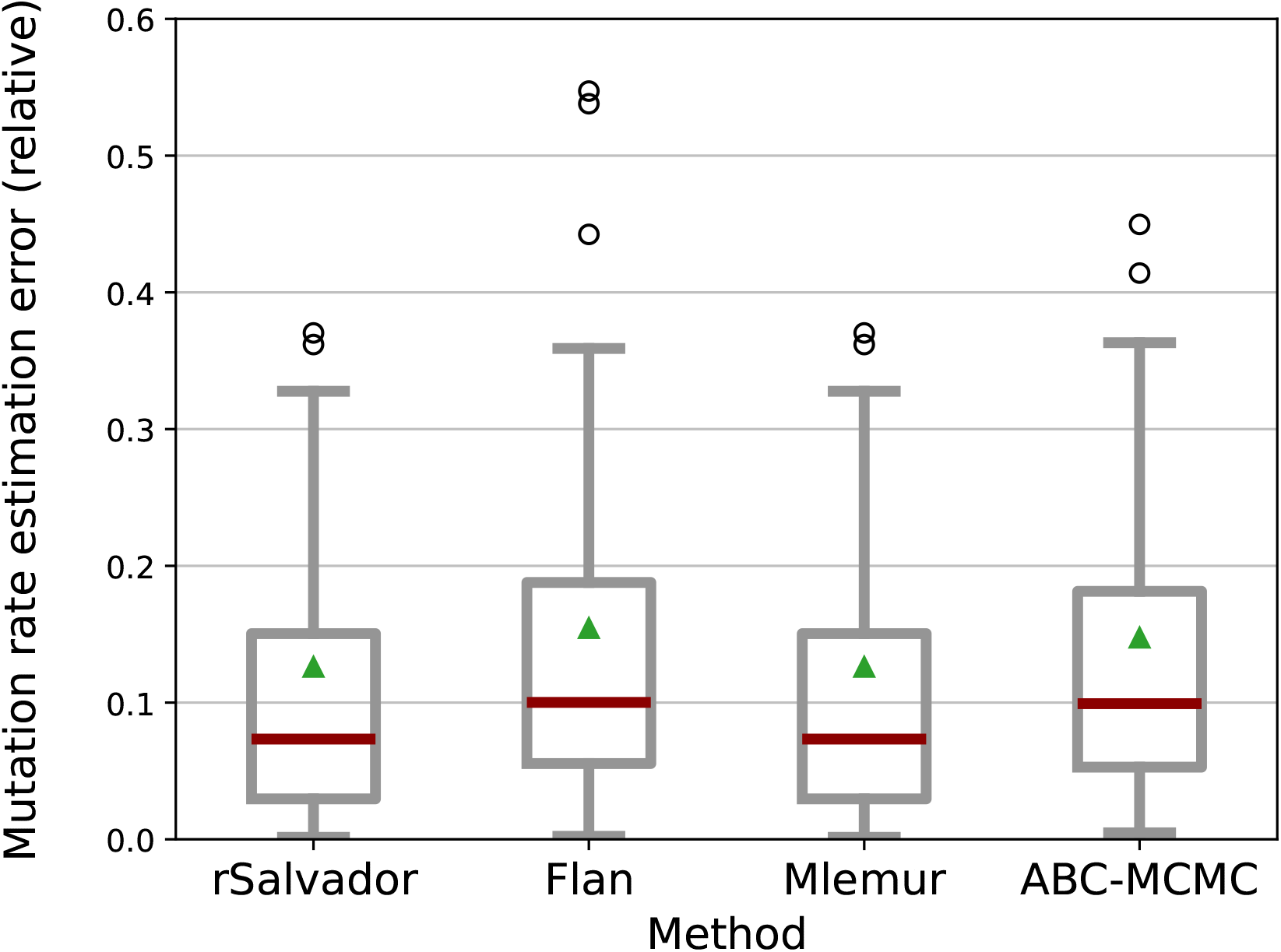
Mutation rate estimation in standard conditions, using rSalvador simulated data. Mutation rate (*µ*) estimation errors through rSalvador, Flan, mlemur, and ABC-MCMC (our proposed method). We performed 160 simulations of fluctuation assays with 24 replicate populations (*r*) for each. For each simulated data, *µ* was randomly drawn from a log-uniform distribution over the range [10^*−*10^, 10^*−*6^], *N* was subsequently randomly drawn from a log-uniform distribution over the range [10^*−*2^, 10^2^]*/µ. µ* is the only unknown parameter, other parameters were set to match the standard assumptions: no fitness difference between mutants and wild types (*f* =1), no death (*d*=0), full population plated without sampling and every mutant detected (*p*=1). Medians are shown with red lines, means are shown with green triangles, the grey boxes represent the 25th and 75th percentile.

